# *Toxoplasma* scavenges mammalian host organelles through usurpation of host ESCRT-III and Vps4

**DOI:** 10.1101/2022.04.11.487923

**Authors:** Julia D. Romano, Joshua Mayoral, Rebekah B. Guevara, Yolanda Rivera-Cuevas, Vern B. Carruthers, Louis M. Weiss, Isabelle Coppens

## Abstract

Intracellular pathogens exploit cellular resources through host cell manipulation. Within its nonfusogenic parasitophorous vacuole (PV), *Toxoplasma* targets host nutrient-filled organelles and sequesters them into the PV through deep invaginations of the PV membrane (PVM) that ultimately detach from this membrane. Some of these invaginations are generated by an intravacuolar network (IVN) of parasite-derived tubules fusing with the PVM. Here, we examine the parasite usurpation of host ESCRT-III and Vps4 to create PVM buds and vesicles. CHMP4B associates with the PVM/IVN and dominant negative (DN) CHMP4B forms many long PVM invaginations containing CHMP4B filaments; the invaginations are shorter in IVN-deficient parasites, suggesting cooperation between IVN and ESCRT. In infected cells expressing Vps4-DN, enlarged intra-PV structures containing host endo-lysosomes accumulate, reflecting defects in PVM scission. Parasite mutants lacking TgGRA14 or TgGRA64 that interact with ESCRT have reduced CHMP4B-DN-induced PVM invaginations and intra-PV host organelles, with greater defects in a double-knockout, revealing the exploitation of ESCRT to scavenge host organelles by *Toxoplasma*.

**Summary:** The parasite *Toxoplasma* sequesters host nutrient-filled organelles into its parasitophorous vacuole through its exploitation of host ESCRT-III and Vps4 for vacuolar membrane-remodeling and fission processes utilizing the parasite proteins TgGRA14 and TgGRA64 that interact with ESCRT.

## Introduction

Biotrophic microbes adapt to life inside host cells by evading host responses to infection and by exploiting surrounding resources to promote multiplication. The apicomplexan protozoan *Toxoplasma gondii* is an obligate intracellular parasite that invades mammalian cells and creates its own replicative niche, the parasitophorous vacuole (PV) in the host cell cytoplasm. Found worldwide, *Toxoplasma* is one of the most successful parasites, capable of reproducing within almost all nucleated cells of warm-blooded animals including humans (Webster, 2010). One critical factor of the success of *Toxoplasma* post-invasion (p.i.) is the secretion of unique proteins from parasite dense granule organelles that transform the PV into a replicative niche and modify the host cell. These proteins, usually named “GRAs” (64 in number so far) exert broad functions ranging from nutrient uptake, host organelle attraction and protein export from the PV into the host cell to control cellular signaling pathways, in particular those involved in innate immune responses and cell cycle (reviewed in (Mercier and Cesbron-Delauw, 2015; Panas and Boothroyd, 2021). For example, TgGRA17 and TgGRA23 form a channel in the PV membrane (PVM) for the import/export of small molecules (<1.9 kDa) (Schwab et al., 1994; Gold et al., 2015). At the PVM, TgMAF1 mediates the association of host mitochondria with the PV (Pernas et al., 2014) while TgGRA3 forms homodimers that govern the trafficking of host Golgi vesicles (Deffieu et al., 2019) that concentrate around the PV to facilitate sphingolipid uptake by *Toxoplasma* (Melo and Souza, 1996; Romano et al., 2013). TgMYR1, TgMYR2 and TgMYR3 form a PVM complex with TgMYR4, TgGRA44/45, and ROP17 to export GRAs into the host cell (Marino et al., 2018; Franco et al., 2016; Cygan et al., 2020; Blakely et al., 2020; Wang et al., 2020) to modify cellular pathways and transcription (reviewed in (Panas and Boothroyd, 2021)).

Within *Toxoplasma*-infected cells, many cellular structures are reorganized and intercepted by the parasite. The host microtubular network is rearranged around the PV, with the host microtubule-organizing center relocating to the PV from the host nucleus (Melo et al., 2001; Coppens et al., 2006; Walker et al., 2008; Romano et al., 2008), allowing host organelles translocating along microtubules to cluster near the PV. Some microtubules create invaginations of the PVM in which host organelles are engulfed (Coppens et al., 2006); blocking microtubule function by treatment of infected cells with Taxol interferes with the transport and sequestration of host endo-lysosomes in the PV. Inside the PV, membranous tubules secreted ‘en bloc’ by the parasite form an elaborate intravacuolar network (IVN) that spreads out in the PV lumen and is maintained by two tubulogenic proteins, TgGRA2 and TgGRA6 (Sibley et al., 1995; Mercier et al., 2002; Bittame et al., 2015). Some IVN tubules connect to the PVM (de Souza and Attias, 2015; Magno et al., 2005) and form open conduits entrapping host organelles, resulting in their sequestration into the PV (Coppens et al., 2006; Romano et al., 2017; Hartman et al., 2022); *Toxoplasma* mutants with a defective IVN (Δ*gra2* and Δ*gra2*Δ*gra6*) have a reduced number of intra-PV Rab vesicles and endo-lysosomes. *Toxoplasma* secretes a phospholipase TgLCAT into the PV that localizes to IVN membranes surrounding internalized host organelles. TgLCAT destabilizes membranes by producing lysophospholipids from the release of fatty acids from phospholipids (Pszenny et al., 2016). Overexpression of TgLCAT leads to fewer intra-PV Rab vesicles, suggesting that TgLCAT hydrolyzes IVN-trapped host organelles, potentially releasing their content into the PV for availability to the parasite.

Our observations by microscopy illustrate that host organelles initially trapped in PVM invaginations (formed by host microtubules or IVN-derived) accumulate in the middle of the PV, suggesting their dissociation from the PVM (Romano et al., 2017). Tubule detachment from the PVM may be analogous to vesicle formation within endosomal compartments, such as the biogenesis of intralumenal vesicles (ILV) from budded portions of the limiting membrane of multivesicular bodies (MVB). This process of membrane invagination and vesicle scission is mediated by Endosomal Sorting Complex Required for Transport (ESCRT) components, with the ESCRT-III complex and the associated AAA-ATPase Vps4 responsible for membrane scission (Piper and Katzmann, 2007; Schöneberg et al., 2017). While some endocytic (e.g., dynamin) and ESCRT components are expressed in *Toxoplasma*, none are secreted into the PV (Nevin and Dacks, 2009; Jimenez-Ruiz et al., 2016; Leung et al., 2008).

The core ESCRT machinery can be subdivided into distinct subcomplexes (ESCRT-I, ESCRT-II, ESCRT-III) and accessory proteins, e.g., apoptosis linked gene 2 (ALG-2, also known as PDCD6) and ALG-2-interacting protein X (ALIX) (Vietri et al., 2020; Schöneberg et al., 2017). ESCRT-III directly reshapes and severs membranes in conjunction with Vps4 while the other subcomplexes and accessory proteins are involved in targeting and activating ESCRT-III (Vietri et al., 2020; Schöneberg et al., 2017; Pfitzner et al., 2021). ESCRT-III is composed of oligomers of Charged Multivesicular Body Protein (CHMP), e.g., CHMP2A/B, CHMP3, CHMP4A/B, with CHMP4 as the most abundant component (Teis et al., 2008; Schöneberg et al., 2017; Vietri et al., 2020). Through an auto-inhibition domain, CHMP proteins form inactive monomers in the cytosol, and upon release of auto-inhibition, activated CHMP proteins polymerize into filaments on membranes and adopt a variety of secondary shapes, resulting in membrane remodeling and scission (Alonso Y Adell et al., 2016; Cashikar et al., 2014; Harker-Kirschneck et al., 2019).

Association of host ESCRT components with the PV of *Toxoplasma* has recently become an active area of research. *Toxoplasma* recruits and utilizes host ALIX and tumor susceptibility gene 101 protein (TSG101) from ESCRT-I during parasite invasion (Guérin et al., 2017). A proximity-labeling study on *Toxoplasma*-infected fibroblasts to identify host proteins at the PV, reveals that a PVM reporter biotinylates several host ESCRT components (e.g., ALIX, ALG-2, TSG101, VPS28, CHMP4B, CC2D1A) (Cygan et al., 2021). Two concomitant studies using co-immunoprecipitation and proximity-based biotinylation identified two *Toxoplasma* proteins on the PVM/IVN that interact with host ESCRT: TgGRA14 (e.g., ALIX, ALG-2, TSG101, CHMP4A/B) (Rivera-Cuevas et al., 2021) and TgGRA64 (e.g., TSG101, VPS37A, VPS28, CHMP4B, ALG-2) (Mayoral et al., 2022). Some of these host ESCRT components have been shown to associate with the PV; ALIX and ALG-2 localize as puncta on the PV (Cygan et al., 2021) and TgGRA14 influences PV recruitment of TSG101 (Rivera-Cuevas et al., 2021). The uptake of host cytosolic proteins by *Toxoplasma* (Dou et al., 2014) is reduced in mammalian cells ectopically expressing the Vps4A EQ dominant negative (DN) mutant or by Δ*gra14* mutant parasites (Rivera-Cuevas et al., 2021).

Based on this information, we hypothesize that host ESCRT machinery diverted by *Toxoplasma* is involved in the scission of IVN tubules containing host vesicles from the PVM. In this study, we analyzed by fluorescence microscopy and transmission EM (TEM) the recruitment of the ESCRT-III component CHMP4B and Vps4 to the PV to assess their impact on PVM remodeling for host organelle internalization. We also examined the distribution of several host ESCRT components on the IVN where intra-PV host organelles accumulate. Finally, we compared the ability of wild-type (WT) and mutants lacking *gra14* and/or *gra64* to scavenge host Rab11 vesicles.

## Results

### Endogenous ESCRT-III component CHMP4B is recruited to the *Toxoplasma* PV and partially associates with host Rab11A vesicles on the IVN

A principle target of intravacuolar *Toxoplasma* is the endosomal recycling pathway in the host mammalian cell (Romano et al., 2017; Hartman et al., 2022). Rab11A localizes to the endocytic recycling compartment (ERC)/recycling endosome and trans-Golgi network to regulate vesicular trafficking through and from these compartments (Maxfield and McGraw, 2004). Endogenous Rab11A in uninfected HeLa cells localized as puncta throughout the cytoplasm with a concentration near the nucleus where the ERC is predominantly located (Fig. 1 A). In *Toxoplasma*-infected cells, the PV is localized close to the host ERC and host Rab11A-positive vesicles. *Toxoplasma* internalizes host GFP-Rab11A vesicles into the PV (89%) using IVN tubules that are appended to the PVM, creating conduits into the PV (Coppens et al., 2006; Romano et al., 2017). We detected several endogenous Rab11A foci on IVN patches (TgGRA7) (Fig. 1A), as previously reported (Romano et al., 2017). The exploitation of IVN tubules to sequester host organelles was further confirmed by TEM of infected cells incubated with LDL-gold particles to track host endo-lysosomes. Data illustrate IVN tubules attached to the PVM (Fig. 1 B, panel a), with some containing LDL-containing organelles in the process of penetrating into a tubule lumen (Fig. 1 B, panel b) or inside a tubule distant from the PVM (Fig. 1 B, panel c). To investigate the hypothesis that host ESCRT machinery is recruited by *Toxoplasma* to induce PVM invaginations and buds into the PV lumen to facilitate host organelle internalization via the IVN, we first monitored the distribution of endogenous CHMP4B upon *Toxoplasma* infection. CHMP4B was detected throughout the cytoplasm in uninfected HeLa cells but, in infected cells, CHMP4B was observed as discrete puncta along the edge of the PV (identified with antibodies against the PVM/IVN protein TgGRA7) and as intra-PV puncta where the IVN concentrates (Fig. 1 C), suggestive of recruitment.

**Figure 1.**
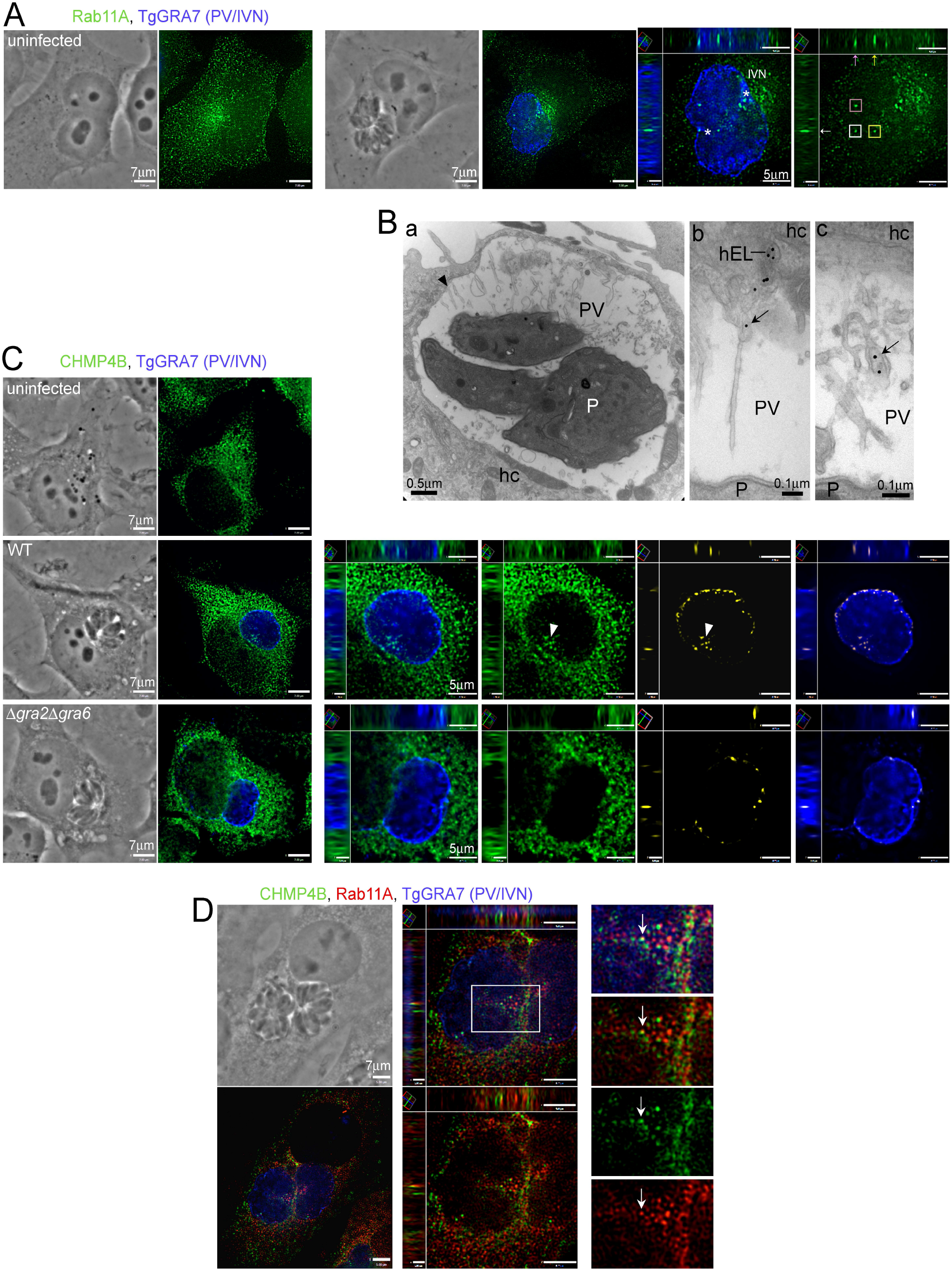
Localization of mammalian CHMP4B and Rab11A in *Toxoplasma*-infected cells. (A) IFA. Uninfected and HeLa cells infected with *Toxoplasma* for 20 h were fixed and immunostained for Rab11A and TgGRA7 (*Toxoplasma* PVM/IVN protein). Individual z-slices and cropped images of the PV (orthogonal views) are shown. Intra-PV Rab11A puncta are shown in squares. Asterisks show patches of IVN. (B) EM. *Toxoplasma*-infected VERO cells were incubated with LDL-gold particles for 24 h before fixation. Arrowhead shows an IVN tubule attached to the PVM. hc, host cell; hEL, host endo-lysosome; P, parasite. (C) IFA. Uninfected and HeLa cells were infected with parasites (WT and Δ*gra2*Δ*gra6*) for 20 h, fixed and immunostained for CHMP4B and TgGRA7. A positive product of the difference from the mean (PDM) image is shown, highlighting in yellow the voxels where the intensity values of CHMP4B and TgGRA7 are above their respective means (overlapping signal), thresholded for background. Individual z-slices and cropped images of the PV (orthogonal views) are shown. Arrowheads show intra-PV CHMP4B puncta. (D) IFA. VERO cells expressing GFP-Rab11A (pseudocolored in red) cells infected with *Toxoplasma* for 20 h were fixed and immunostained CHMP4B and TgGRA7. Individual z-slices and cropped images of the PV (orthogonal views) are shown. Arrows point to an overlapping CHMP4B and Rab11A puncta on the IVN.

To further examine the possible role of the IVN in host CHMP4B delivery into the PV, we monitored endogenous CHMP4B in HeLa cells infected with the Δ*gra2*Δ*gra6* mutant that has a defective IVN (Mercier et al., 2002) and is deficient in host vesicle internalization (Romano et al., 2017). CHMP4B was also observed as puncta along the periphery of the Δ*gra2*Δ*gra6* PV, but little to no puncta were detected in the PV lumen (Fig. 1 C), suggesting a contribution of IVN tubules to host CHMP4B entry into the PV. We then assessed the intra-PV distribution of host CHMP4B relative to Rab11A vesicles trapped in the IVN; a partial overlap was observed between CHMP4B and Rab11A puncta (Fig. 1 D), suggesting a possible involvement of host CHMP4B in host vesicle internalization into the PV in synergy with IVN tubules.

### Dominant negative CHMP4B tagged with mEmerald at the N- or C-terminus is targeted to different locations at the PV

Host CHMP4B partially localizes to the PVM and the IVN, supporting the hypothesis that *T. gondii* utilizes the host ESCRT-III complex during infection. CHMP4B undergoes a conformational change between a closed, soluble and inactive form and an open, active conformation that exposes C-terminal domains to recruit Vps4 and other effectors, such as ALIX, enabling membrane binding and oligomerization (Tang et al., 2015; Muziol et al., 2006; Lata et al., 2008; Obita et al., 2007; McCullough et al., 2008; Henne et al., 2012; Lin et al., 2005). Ectopic expression of CHMP proteins with bulky protein tags on either the N- or C-terminus favors the open conformation and causes DN effects leading to the formation of enlarged endosomal “class E” compartments (Martin-Serrano et al., 2003; Howard et al., 2001; von Schwedler et al., 2003). Taking advantage of the DN effect that ESCRT-III subunits tagged with large fluorophores have on complex activity, we next investigated the DN effect of CHMP4B on the PV. We expressed CHMP4B tagged with mEmerald either at the N-(mEm-CHMP4B) or C-(CHMP4B-mEm) terminus in HeLa cells. In uninfected cells expressing either construct, large punctate structures consistent with enlarged “class E” compartments containing tagged CHMP4B were observed throughout the cytoplasm (Fig. 2, A and B). In cells expressing mEm-CHMP4B and infected with WT parasites for 20 h, mammalian “class E” compartments surrounded the PV and discrete mEM-CHMP4B puncta were detected on the PVM and also inside the PV within IVN patches (Fig. 2 A, Fig. S1 A). In cells infected with the IVN mutant Δ*gra2*Δ*gra6*, the mEm-CHMP4B compartment associated with the PV but no puncta were observed within; faint mEm-CHMP4B staining was observed along the PVM (Fig. 2 A). In contrast, in infected cells expressing CHMP4B-mEmerald, the signal was predominantly distributed on the PVM of both WT and Δ*gra2*Δ*gra6* parasites, and no puncta were observed within the PV (Fig. 2 B, Fig. S1 B).

**Figure 2.**
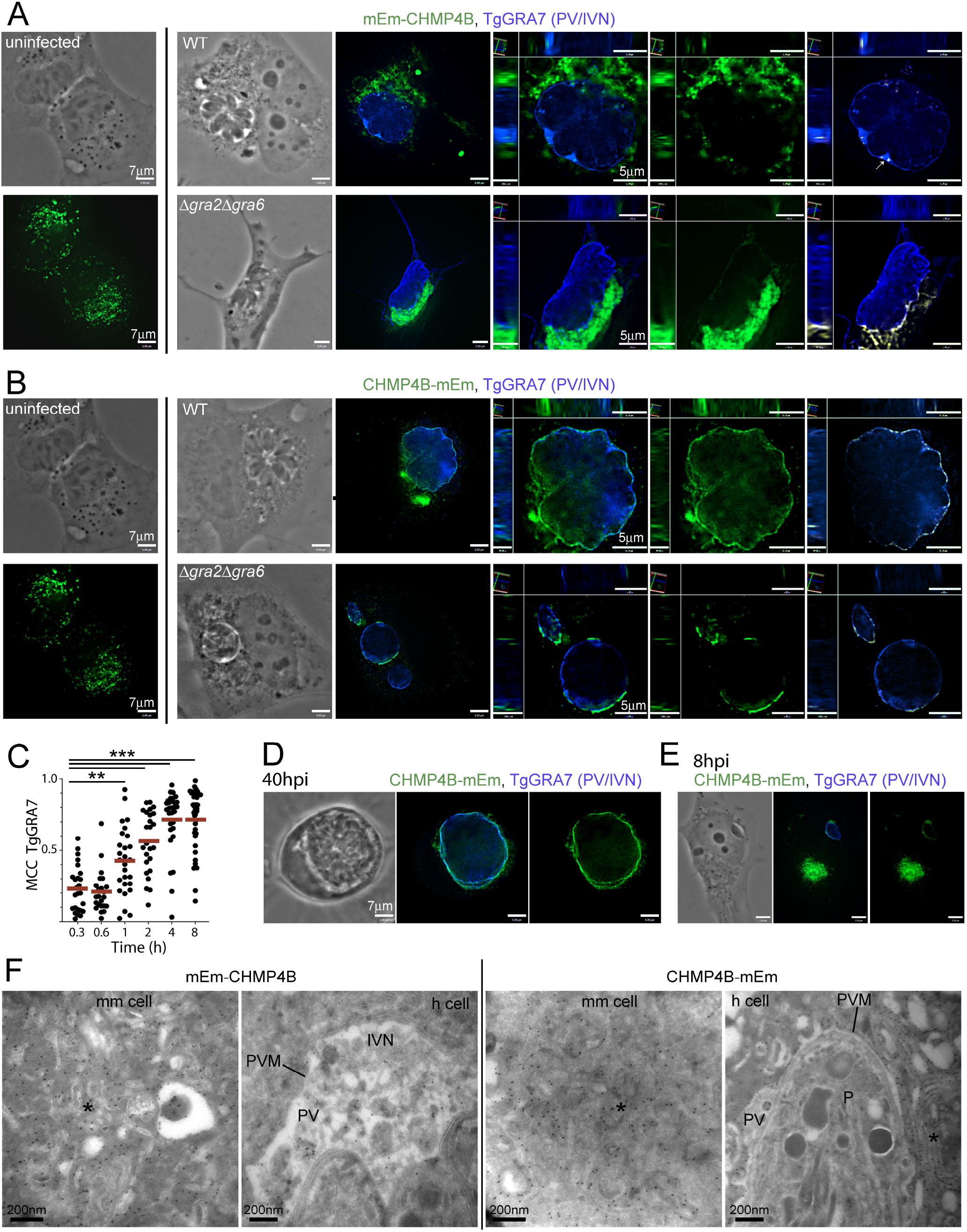
Differential PV localization of CHMP4B tagged with either N- or C-terminally mEmerald. (A-B) IFA. HeLa cells were transiently transfected with either mEm-CHMP4B (A) or CHMP4B-mEm (B). Uninfected and cells infected with parasites (WT or Δ*gra2*Δ*gra6*) for 20 h were fixed and immunostained for TgGRA7. Arrow in A shows CHMP4B staining on the IVN. Individual z-slices and cropped images of the PV (orthogonal views) are shown plus positive PDM images. (C) IFA. HeLa cells transiently transfected with CHMP4B-mEm were infected with WT parasites at the indicated times, fixed and immunostained for TgGRA7. Mander’s correlation coefficients (MCC, TgGRA7) were calculated with Volocity. Dot blots are shown with red bars indicating the mean. At least 20 PV were measured per time point. p-values, ***<0.0001, **=0.0098. (D-E) IFA. HeLa transiently transfected with CHMP4B-mEm were infected with WT parasites for 40 h (D) or 8 h (E), fixed and immunostained for TgGRA7. Individual z-slices are shown. (F-G) ImmunoEM. HeLa cells transiently transfected with either mEm-CHMP4B (F) or CHMP4B-mEm (G) and either uninfected (left panels) or infected with WT parasites for 20 h. Asterisks show “Class E”-like compartments. mm cell, mammalian cell; h cell, host cell; P, parasite.

We next wanted to monitor the recruitment of CHMP4B-mEm at the PV over time by measuring the levels of colocalization of CHMP4B-mEm and TgGRA7 at the PVM using the Mander’s correlation coefficient (MCC). We observed that CHMP4B-mEm progressively associated with the PVM (Fig. S1 C), with a mean MCC value for TgGRA7 of 0.232 at 20 min p.i. and increasing to 0.716 at 8 h p.i. (a time point typically corresponding to the first round of *Toxoplasma* division) (Fig. 2 C, Fig. S1 C). The mean MCC values for CHMP4B-mEm also trended higher as the infection progresses, though not all of CHMP4B-mEm associated with the PVM at 8 h p.i. (Fig. S1 C and D). However, late in infection at 40 h p.i. prior to *Toxoplasma* egress, CHMP4B-mEm staining was mainly detected on the PVM, suggesting continuous recruitment by the parasite (Fig. 2 D). Interestingly, CHMP4B association with the PV occurred even when the bulk of CHMP4B-mEm localized far away from the PV, e.g., on the other side of the host nucleus (Fig. 2 E) and was also observed along projections of the PVM that extend into the host cytosol (Fig. S1 C, 8 h p.i.).

To scrutinize the association of mEmerald-tagged CHMP4B with the PVM in finer detail, we performed immuno-EM using anti-GFP antibodies on infected cells expressing either construct. Within the mammalian cell, gold particles localized mEm-CHMP4B and CHMP4B-mEm to multilamellar “class E” compartments (Fig. 2 F). In infected cells, PV-associated mEm-CHMP4B predominantly distributed to IVN tubules while gold particles detecting CHMP4B-mEm were observed aligned along the PVM, covering the PV entirely on each cryosection. Altogether, these observations highlight that after induction of DN CHMP4B in infected cells, *Toxoplasma* actively recruits CHMP4B to the PV, but the final destination of this protein on the PV differs according to the mEmerald tag position. Our data indicate that mEm-CHMP4B found predominantly on the IVN, is internalized into the PV while CHMP4B-mEm, localized uniformly to the PVM but is absent from the PV interior indicating that CHMP-4B-mEm is blocked at this membrane. This suggests that the C-term mEm tag allows initial insertion of CHMP4B to the PVM but interferes with later steps in scission from this membrane.

### The PVM containing host CHMP4B-mEmerald invaginates and buds into the PV lumen

The massive recruitment of CHMP4B-mEm at the PVM prompts us to examine in great detail and resolution the ultrastructure of this membrane in ultra-thin resin embedded cells. In non-transfected cells, the PVM had a relatively flat appearance with closely apposed host mitochondria and ER elements (Fig. 3 A, panel a). These host organelles form intimate association with the PVM mediated by parasite proteins (Melo et al., 1992; Melo and Souza, 1997; Sinai et al., 1997; Pernas and Boothroyd, 2010; Pernas et al., 2014) but are never detected within the PV, unlike host endocytic organelles. Higher magnification observations reveal PVM indentations containing host LDL-containing organelles, seemingly targeted by tubules of the IVN as a point of anchorage and fusion for host organelle internalization into the PV (Fig. 3 A, panel b). Infected cells transfected with CHMP4B-mEm were identified by the presence of “class E” compartments in the host cytosol. Unlike PV in non-transfected cells, the PVM containing CHMP4B-mEm displayed a corrugated morphology, with tubular (up to 450 nm-long) and numerous invaginations (often spaced less than ∼50 nm spaced from each other) detected over the entire surface of this membrane (Fig. 3 B). In transfected cells expressing CHMP4B-mEm and infected with Δ*gra2*Δ*gra6* mutant parasites, the PVM of the mutant also contained CHMP4B-mEm, though to a lesser extent than WT PVM (Fig. 1 C), and underwent the same deformation process with the generation of nanosized buds away from the host cytoplasm (Fig. 3 C). However, the PVM buds in the Δ*gra2*Δ*gra6* mutant were rarer, further apart and smaller in size (∼80 nm long). This suggests a contribution of IVN tubules in supplying membranes at sites of PVM buds in WT PV.

**Figure 3.**
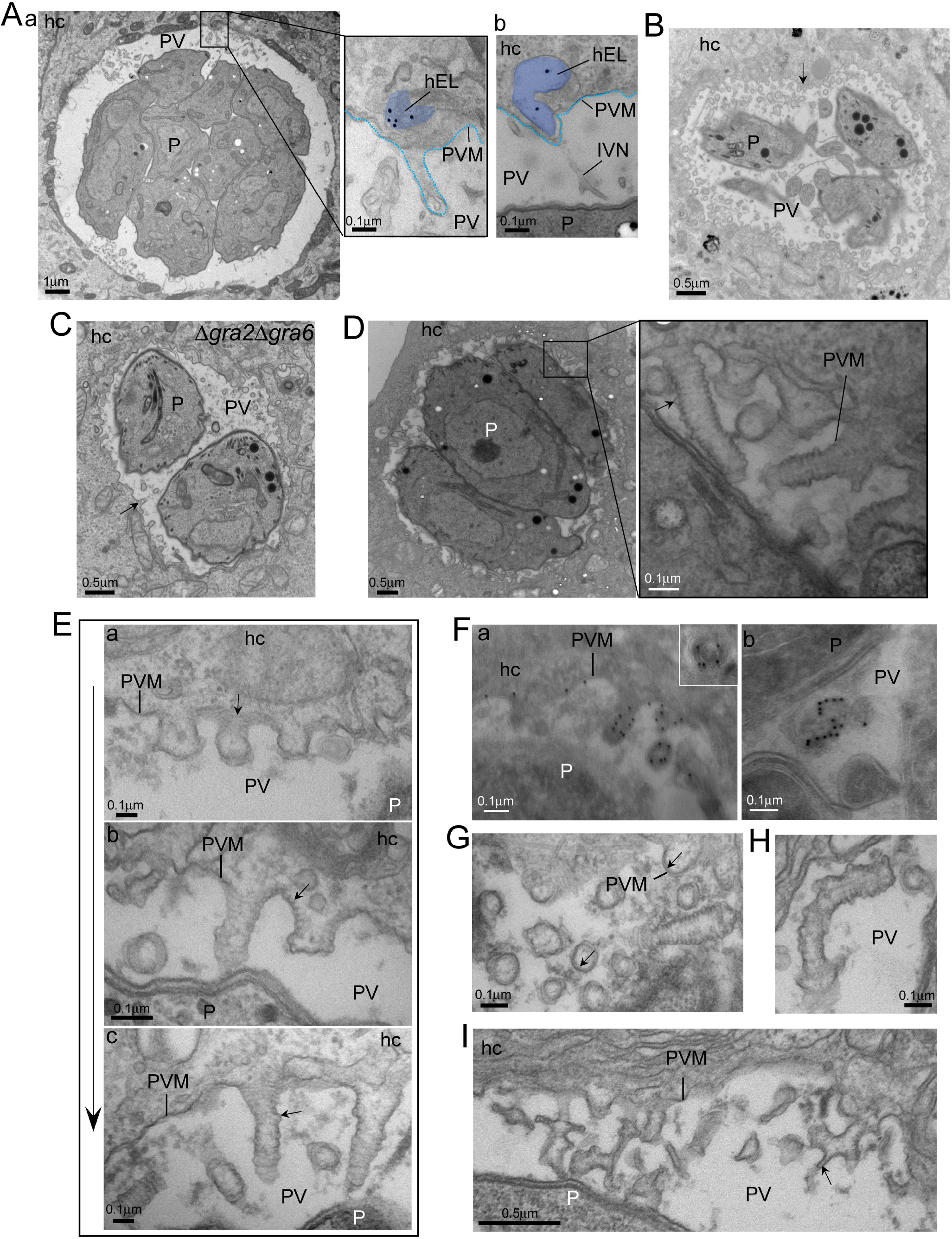
Ultrastructure of PV in HeLa cells expressing CHMP4B-mEmerald. (A-I) EM. (A) HeLa cells infected with *Toxoplasma* were incubated with LDL-gold particles for 20 h to track host endo-lysosomes (hEL, pseudocolored in blue in inset in panels a and b). (B-C) HeLa cells transfected with CHMP4B-mEm and infected with WT parasites (B) showing extensive PVM remodeling or with Δ*gra2*Δ*gra6* parasites (C) showing fewer and smaller tubules at the PVM. Arrows show PVM buds or tubules. (D-I) CHMP4B-mEm-expressing HeLa cells infected with WT parasites detailing numerous large tubular structures showing regular constrictions (arrow in D). In E, panels a to c show examples of different PVM tubule lengths (arrows). In F, immunogold staining with anti-GFP antibody showing gold particles within tubules. In G, transversal sections of tubules showing CHMP4B filaments inside tubules (arrows) identifiable by their higher electron-density compared to tubular membrane. In H and I, illustration of curled or ramified tubules. Arrow in I shows a smaller tubule, likely from the IVN, attached to a large constricted tubule. hc, host cell; P, parasite.

In control cells, intravacuolar *Toxoplasma* distribute radially, forming a rosette-like structure. Consequent to the extensive PVM remodeling upon host CHMP4B-mEm recruitment, the PV occasionally contained spatially disorganized parasites and very few host mitochondria and ER were observed at the PVM (Fig. S2 A). Furthermore, some PV dually stained for CHMP4B and TgGRA7 at the PVM look ‘empty’ as a sign of *Toxoplasma* progressive deterioration (Fig. S2 B and C). In cells expressing CHMP4B-mEm, enumeration of parasites per PV at 20 h p.i. shows that ∼35% of PV contained 8 parasites similarly to cells expressing mEm-CHMP4B, but uniquely ∼15% of PV did not harbor any parasites (Fig. S2 D). Compared to mock transfection, no PV with 16 parasites were observed in cells expressing mEm-CHMP4B or CHMP4B-mEM.

High magnification observations of PVM containing host CHMP4B-mEmerald reveal, on longitudinal sections of tubular invaginations, regularly displayed constrictions on a horizontal plan and continuing over the length of the tubular structures, resulting in a reduction of the membrane tube radius (Fig. 3 D). The overall diameter of the PVM tubules was uniform, corresponding to 105 ± 10 nm. Our morphological analyses suggest sequential steps in the formation of PVM tubular invaginations. Initially, distinct filamentous structures, likely CHMP4B polymers, were observed accumulated at many areas of the PVM. Then, these confined areas decorated with filaments evolved from flat to small buds (Fig. 3 E, panel a). Filaments were detected in the interior of the buds, organized into rings attached to membrane (Fig. 3 E, panel b). Finally, concomitant to the incorporation of filaments into PVM buds, these latter transformed progressively into long tubules up to 8 µm that underwent constriction with decreased diameter (Fig. 3 E, panel c). Immunostaining for CHMP4B confirms the presence of gold particles coating the interior of PVM invaginations (Fig. 3 F, panel a). Stabilized ESCRT-III filaments including CHMP4B polymers, are frequently organized into spirals on curved membranes (Cashikar et al., 2014). A similar organization of CHMP4B filaments was observed within PVM tubules, as observed in a tangential section showing gold particle distribution in a spiraled pattern (S-shape) (Fig. 3 F, panel b) and on transversal sections of PVM tubules showing CHMP4B filaments (characterized by high electron-density layer beneath the tubule membrane) forming a partial ring that may reflect a spiral configuration (Fig. 3 G). We also noticed that some PVM tubules appeared slightly bent (Fig. 3 D and H) and branched, with some bifurcations reminiscent of IVN tubules that may have fused with large PVM tubules containing CHMP4B filaments (Fig. 3 D and I).

Altogether these observations suggest that host CHMP4B recruited at the PVM assemble into filaments at the surface of this membrane, driving tubular invaginations away from the cytoplasm. Incarcerated within PVM tubules, assembled CHMP4B filaments narrow the tubule, similar to the CHMP4A internal coat on negatively-curved membranes to draw membranes together to the fission point (McCullough et al., 2015).

### FLAG-tagged CHMP4B localizes to the PVM and as enlarged puncta within the PV close to Rab11A vesicles

Overexpression of CHMP proteins fused to a protein tag leads to DN effects, presumably due to the bulky tag interfering with inhibitory intramolecular interactions necessary to keep CHMP proteins in an inactive soluble form. Because of the strong DN effect of CHMP4B tagged with mEm, we opted to transfect infected cells with CHMP4B in fusion with a FLAG tag (8 amino acids; ∼1 kDa) at the N-terminus (FLAG-CHMP4B). Previous studies reported FLAG-CHMP4B localization as a punctate pattern upon transient transfection in HeLa cells and a more diffuse cytoplasmic distribution upon stable expression in HEK293 cells, with vesicle-like structures observed in 5% of the cells (Katoh et al., 2003). We confirmed the localization of FLAG-CHMP4B as disperse puncta, partially concentrated in the perinuclear region in HeLa cells, as a result of overexpression (Fig. 4 A, panel a). In HeLa cells infected with *T. gondii* for 20 h, three patterns for FLAG-CHMP4B distribution were observed: 66% of cells displayed a FLAG-CHMP4B signal throughout the host cell with partial localization to the PV, while 34% exhibited a signal predominantly associated with the PV (Fig. 4 A, panel b). Among the 34% population, FLAG-CHMP4B was observed either as very large intra-PV puncta (up to 1 µm in diameter) near IVN patches (23%) or all along the PVM (11%). The majority of large intra-PV puncta containing FLAG-CHMP4B had a torus shape that colocalized with TgGRA7, indicating that CHMP4B puncta are derived from the PVM (Fig. 4 A, panel c). To assess if IVN tubules could be involved in the internalization of FLAG-CHMP4B into the PV, FLAG-CHMP4B-expressing cells were infected with Δ*gra2*Δ*gra6* parasites. Two patterns for FLAG-CHMP4B distribution in infected cells were discernible, corresponding to partial staining of the PV with the bulk in the host cytoplasm (58%) or to preponderant PV labeling (42%) but with no intra-PV puncta detected (Fig. 4 A, panel d).

**Figure 4.**
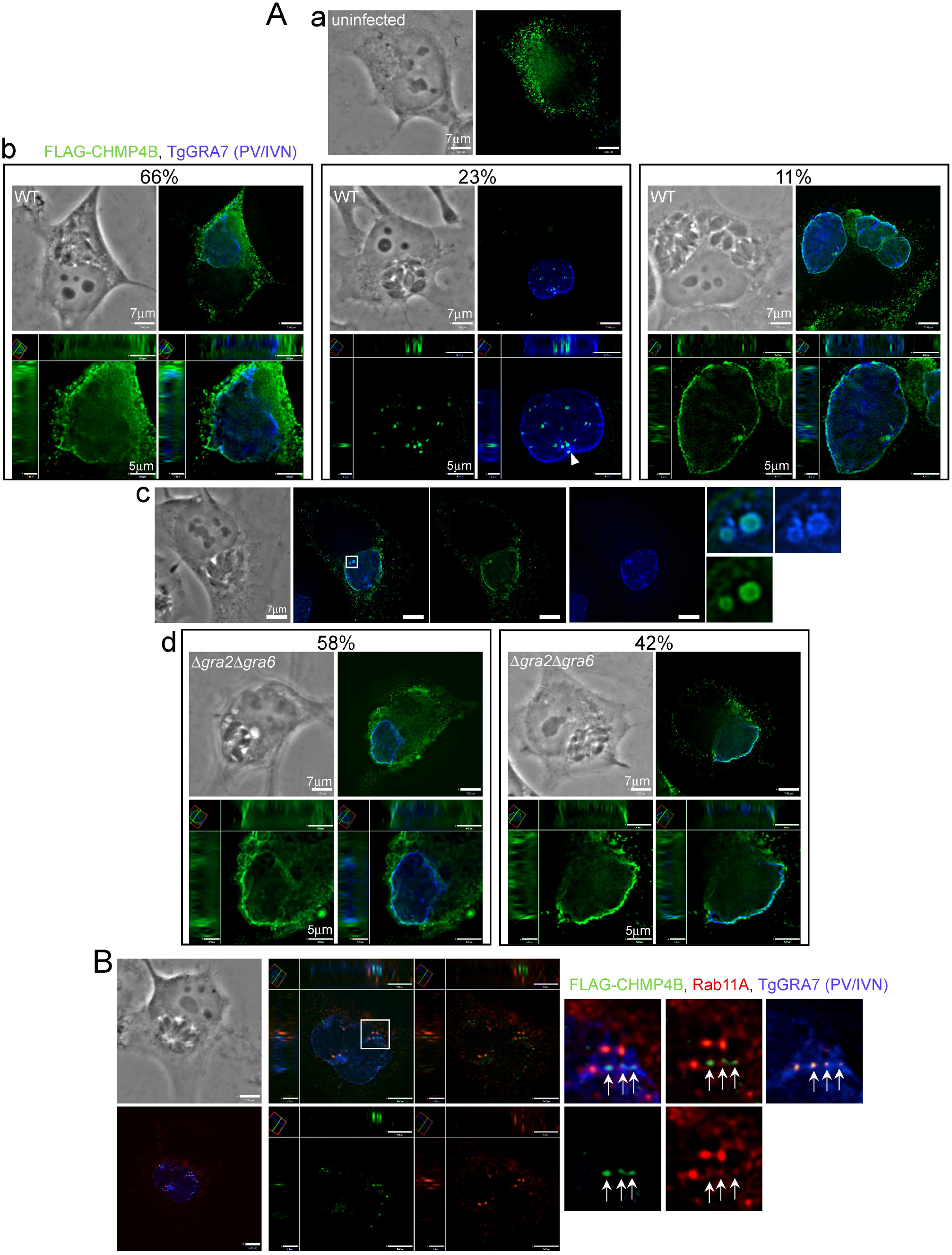
Association of FLAG-CHMP4B with the PV. (A) IFA. HeLa cells were transiently transfected with FLAG-CHMP4B and either uninfected (panel a) or infected with WT parasites (panels b and c) or Δ*gra2*Δ*gra6* parasites (panel d) for 20 h, fixed and immunostained for FLAG and TgGRA7. Shown are individual z-slices and cropped images of the PV (orthogonal views). The percent of infected cells with the displayed morphology is shown. (B) IFA. HeLa cells transiently transfected with FLAG-CHMP4B were infected with WT parasites for 20 h, fixed and immunostained for FLAG, Rab11A and TgGRA7. Individual z-slices and cropped images of the PV (orthogonal views) are shown plus positive PDM images (yellow). Enlarged regions show overlap of FLAG-CHMP4B and Rab11A puncta (arrows).

We previously demonstrated that host CHMP4B inside the PV in close proximity to Rab11A vesicles trapped in the IVN, which suggests the promotion of host vesicle internalization mediated by host CHMP4B and IVN tubules. To confirm such a role for CHMP4B, we infected HeLa cells expressing FLAG-CHMP4B and immunostained for Rab11A. Several intra-PV FLAG-CHMP4B and Rab11A puncta were detected in patches of TgGRA7 signal, presumably overlapping with the IVN, and to a lesser extent with each other (Fig. 4 B and Fig. S3 A and B). We next conducted EM observations to scrutinize the PVM in infected HeLa cells expressing FLAG-CHMP4B. In the cytoplasm of transfected cells overexpressing FLAG-CHMP4B, many vesicles of variable size were generated (Fig. 5 A). In all PV examined in transfected cells, no PVM tubulation or unusual remodeling was observed (Fig. 5 B), similar to the PV in cells expressing mEm-CHMP4B (Fig. 2 F). However, the PV lumen contained several large unique structures (up to 700 nm in diameter) composed of numerous vesicles (20 to 100 nm in diameter) encircled by membrane and close to the IVN, with some attached to the PVM (Fig. 5 B and C). These large multivesicular structures were reminiscent of large CHMP4B puncta surrounded by TgGRA7, detected by IFA (Fig. 4 A, panel c). Based on size and electron-density, we interpret the nature of the luminal vesicles to likely be host endosomal vesicles formed upon CHMP4B overexpression that are subsequently internalized into the PV and surrounded by the PVM.

**Figure 5.**
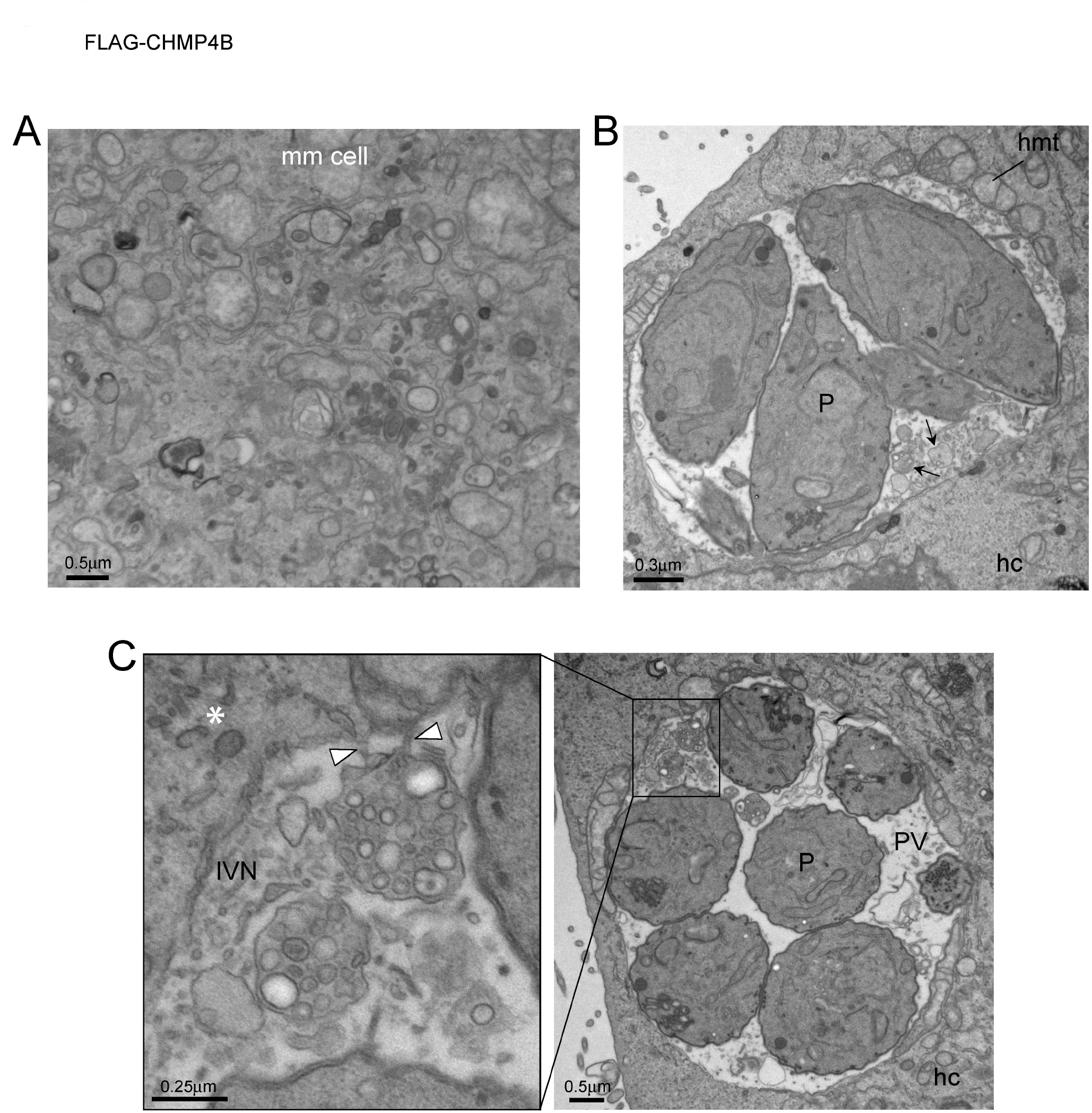
Ultrastructure of infected HeLa cells expressing FLAG-CHMP4B. (A-C) EM. Infected HeLa cells expressing FLAG-CHMP4B showing accumulation of numerous vesicles in the cytoplasm of host cells (A), no PVM remodeling but accumulation of large intra-PV multivesicular structures (arrows), some with visible connection with the PVM (arrowheads in B-C). mm cell, mammalian cell; hc, host cell; hmt, host mitochondria; P, parasite.

### Expression of dominant negative Vps4A results in PV swelling and accumulation of enlarged intra-PV membrane-bound structures containing host organelles

Vps4A and Vps4B proteins are members of a class of hexameric AAA^+^-ATPases that disassemble protein complexes without degradation. Vps4 enzymes contain an N-terminal MIT domain that binds to the tails of ESCRT-III proteins and ESCRT-III/Vps4 complexes assemble within the necks of membrane tubules and convert the energy of ATP hydrolysis into pulling forces that sever the tubules (Yang et al., 2015). Vps4 is also required for the recycling of ESCRT-III from membrane-bound filaments back to the cytosol (Babst et al., 1998; Bishop and Woodman, 2000; Lin et al., 2005). As Vps4 is involved in many cellular processes that require membrane remodeling and scission, we hypothesize that *Toxoplasma* exploits host Vps4 along with CHMP4B to facilitate PVM budding and scission of IVN tubules containing host organelles from the PVM. We analyzed the distribution of Vps4A in infected cells transfected with mCherry-Vps4A. In both transfected VERO and HeLa cells, we observed a perivacuolar staining for mCherry-Vps4A (Fig. 6 A and B). Mean fluorescence intensities for mCherry-Vps4A and TgGRA7 assessed by PDM^+^ values reveal an association of mCherry-Vps4A at some areas on the PVM. In VERO cells stably expressing GFP-Rab11A or in HeLa cells immunostained for Rab11A, a partial overlap between intra-PV GFP-Rab11A and mCherry-Vps4A foci was observed.

**Figure 6.**
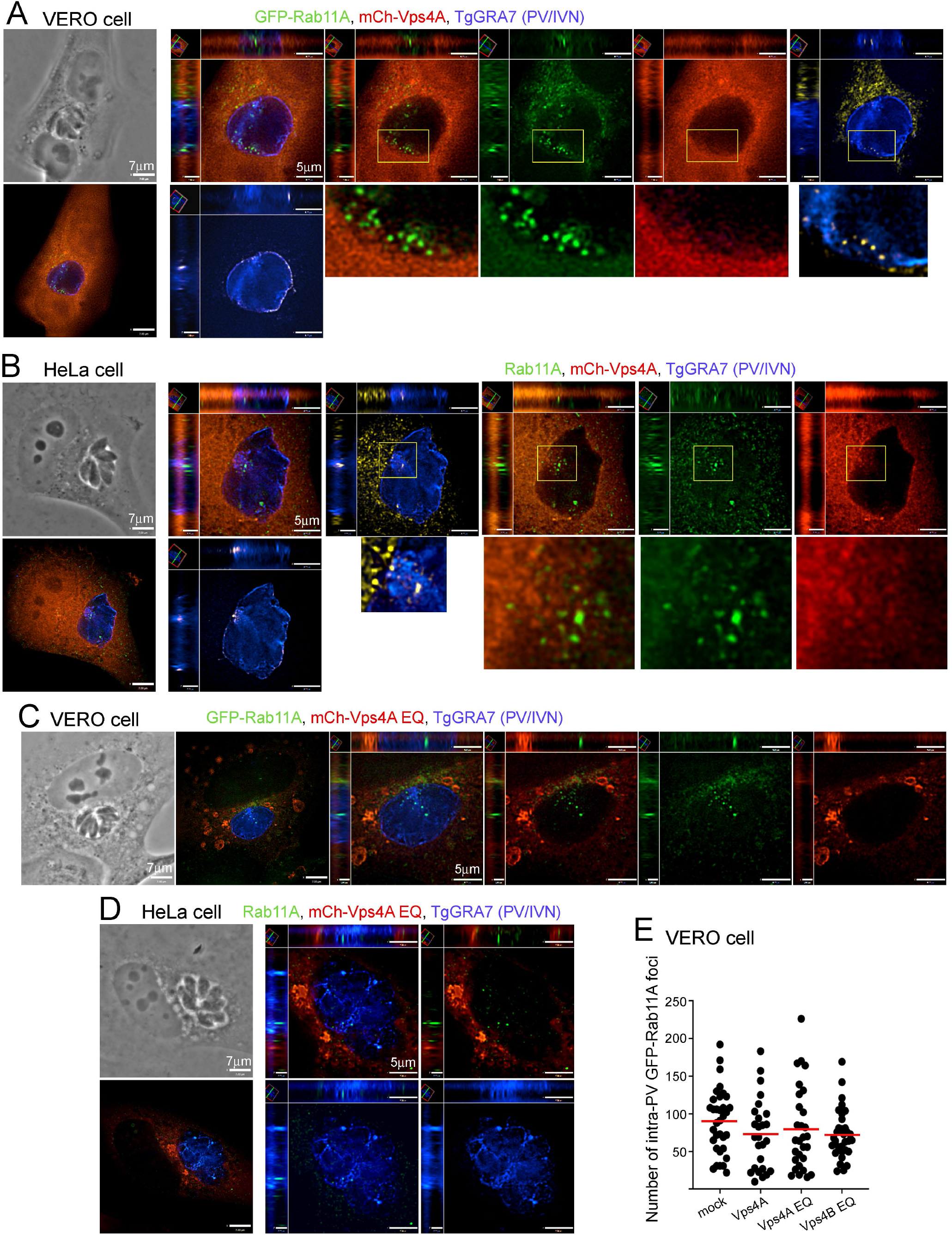
Effect of DN Vps4 on host Rab11A vesicles internalization. (A-D) IFA. VERO cells expressing GFP-Rab11A (A, C) or HeLa cells (B, D) were transiently transfected with either mCherry-Vps4A WT (A, B) or mCherry-Vps4A EQ (DN) (C, D). The cells were infected with WT parasite for 20 h, fixed and immunostained for TgGRA7 (A-D) or Rab11A (B, D). Individual z-slices and cropped images of the PV (orthogonal views) and enlargements of boxed regions are shown plus positive PDM images (yellow). (E) VERO cells expressing GFP-Rab11A were mock-transfected or transiently transfected with mCherry-Vps4A (WT or EQ) or -Vps4B EQ, infected with WT parasites for 20 h, fixed, and immunostained for TgGRA7. The number of intra-PV GFP-Rab11A foci was measured using Volocity software. Dot plots are shown with red bars indicating the mean from one representative experiment. At least 26 PV were analyzed. No significant differences were found (one-way ANOVA).

Expression of the DN mutant Vps4A EQ (in which the glutamate at position 228 in the ATPase active site is replaced by a glutamine) causes a class E phenotype with an aggregation of MVB due to blockade of ESCRT-dependent ILV formation (Bishop and Woodman, 2000). Next, we examined if dysfunction of ESCRT-III modelled by Vps4A EQ could impact host organelle sequestration into the PV. In VERO cells and HeLa cells transfected with mCherry-Vps4A EQ, enlarged endosomal vacuoles were observed, with some surrounding the PV (Fig. 6 C and D), as observed previously (Rivera-Cuevas et al., 2021), but no signal was detected inside the PV. GFP-Rab11A puncta were observed inside the PV, with no significant difference in number as compared to mock-transfected or cells transfected with Vps4A or DN of Vps4B (Fig. 6 E). However, upon expression of Vps4 EQ, the PVM (delineated by immunostaining with TgGRA7) appeared to be abnormally protruding into the PV lumen (Fig. 6 D), suggesting that structural alterations of this membrane may be occurring, even in the absence of detectable Vps4EQ signal on this membrane.

We conducted TEM analyses on infected HeLa cells transfected with mCherry-Vps4EQ to investigate morphological PVM defects and assess the PV for host organelle sequestration. Transfected cells were incubated with LDL-gold particles and expectedly, we detected gold particles in enlarged multilamellar organelles accumulated in the host cytoplasm (Fig. 7 A), with some of them clustering around the PV (Fig. 7 B). A striking observation was the presence of large membranous whorls derived from the PVM and expanding into the vacuolar space (Fig. 7 B and C); some IVN tubules were seen attached to these enlarged structures (Fig. 7 C, right inset). While LDL-gold-containing organelles were observed within the PV, they amassed in bundles forming very large multivesiculated entities up to 900 nm in diameter as quantified in Fig. 7 E), as compared to the small size of LDL-gold-containing organelles in the PV of non-transfected cells (∼200 nm in diameter) (see examples in Fig. 1 B, panel c and Coppens et al., 2006). These LDL-gold-containing organelles remained in close proximity to the PVM with an average distance of ∼0.25 µm, whereas in WT PV many host organelles were present in the PV center (Fig. 7 E). Altogether, these TEM observations reveal that Vps4A DN expression causes abnormal PVM proliferation with entrapment of host organelles in PVM folds, suggesting retardation of scission activities due to impaired functions of Vps4A.

**Figure 7.**
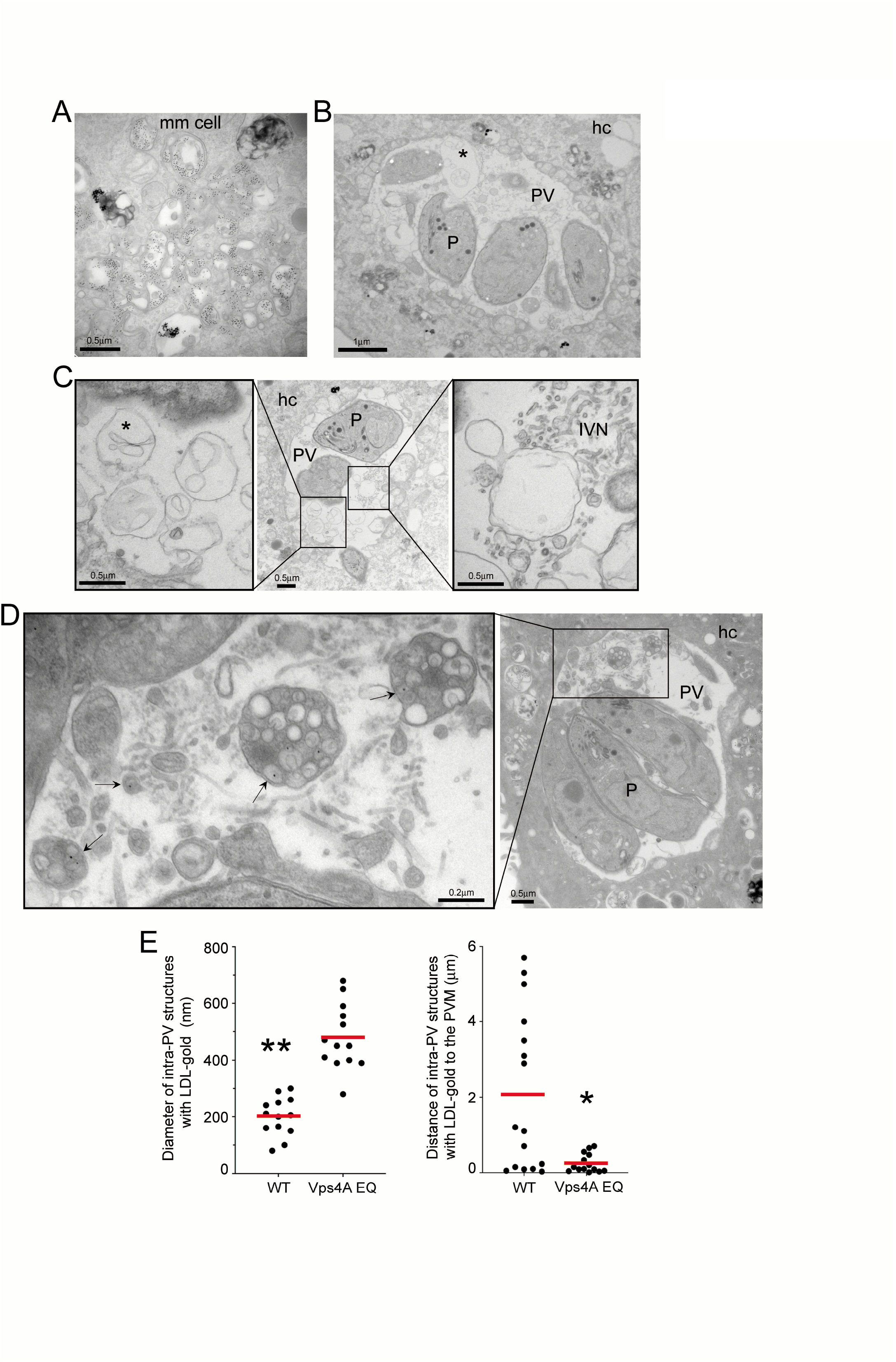
Ultrastructure of PVM and PV content in cells overexpressing Vps4A EQ. (A-D) EM. Infected HeLa cells expressing mCherry-Vps4A EQ and incubated with LDL-gold particles showing class E-like compartments containing gold particles in the host cytoplasm (A) and large membrane buds derived from the PVM (B, C; asterisks), some in contact with IVN tubules shown in C (right inset). In D, intra-PV detection of very large structures with host LDL-gold organelles accumulated (arrows) hc, host cell; P, parasite. (E) Measurement of the size and distance of intra-PV structures containing LDL-gold from their center to the PVM. Dot blots are shown with red bars indicating the mean from one representative experiment. For p-values (unpaired, two-tailed t test), **p<0.0001, *p=0.0032.

### The ESCRT accessory protein ALIX and more predominantly its interacting partner ALG-2 colocalize with intra-PV Rab11A vesicles

ALIX, an adaptor protein whose Bro1 domain binds to a motif at the C-terminus of CHMP4 proteins, participates in a variety of ESCRT-III-mediated membrane remodeling processes (McCullough et al., 2008). ALG-2 is a calcium-binding protein that forms intramolecular interactions with ALIX in a Ca^2+^-dependent manner (Sun et al., 2015). Host ALIX and ALG-2 have been shown previously to associate with the PV of *Toxoplasma*, as puncta along the PVM and within the PV (Cygan et al., 2021; Rivera-Cuevas et al., 2021). To investigate if host ALIX or ALG-2 may participate to host vesicle internalization into the PV, we examined the distribution of intra-PV ALIX or ALG-2 puncta relative to Rab11A vesicles. In infected GFP-Rab11A-expressing VERO cells transfected with HA-tagged ALIX, a few intra-PV ALIX-HA puncta displayed a partial overlap with GFP-Rab11A (Fig. 8 A). In comparison, in infected GFP-Rab11A-expressing VERO cells transfected with ALG-2-HA, the association of ALG-2-HA with the PVM and IVN was more prominent and numerous ALG-2-HA puncta co-localized with GFP-Rab11A (Fig. 8 B). We performed dual immunogold labeling on ALG-2-HA expressing HeLa cells using anti-HA and anti-Rab11A antibodies to assess the colocalization of GFP-Rab11A and ALG-2-HA in finer detail. In the mammalian cytoplasm, gold particles were observed on membranes and vesicles associated with either Rab11A or ALG-2-HA, but none with both (Fig. 8 C, panel a). Within the PV, however, both Rab11A and ALG-2-HA were detected on the same vesicular structures with Rab11A (10-nm-gold particles) on the inside and ALG-2-HA (5-nm-gold particles) externally displayed (Fig. 8 C, panels b-d). This illustrates that intra-PV Rab11A vesicles are surrounded by a membrane presumably derived from the PVM/IVN that contains ALG-2, suggesting a potential role for ALG-2 and to a lesser extent ALIX in cooperating with CHMP4B at sites of PVM remodeling for host organelle sequestration.

**Figure 8.**
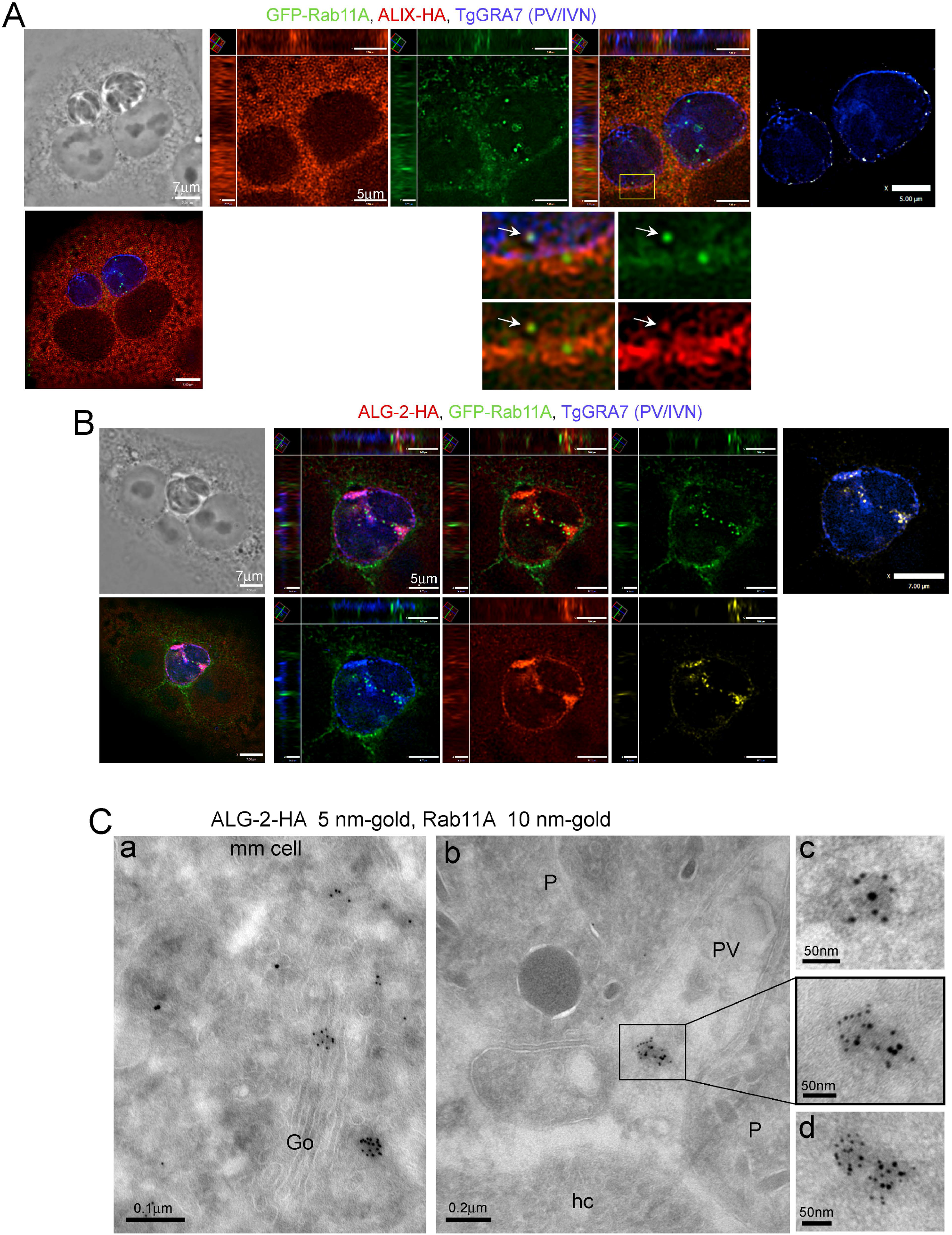
Association of mammalian ALIX and ALG-2 with intra-PV Rab11A vesicles. (A-B). IFA. VERO cells stably expressing GFP-Rab11A were transfected with ALIX-HA (A) or ALG-2-HA (B), infected with WT parasites for 20 h, fixed and immunostained for HA and TgGRA7. Individual z-slices and cropped images of the PV (orthogonal views) are shown. Boxed regions are shown in enlargements and arrows indicate overlap of GFP-Rab11A and ALIX-HA signal (A). In A, positive PDM images correlating ALIX-HA and the PVM (GRA7) (white) and in B, positive PDM images correlating ALGALG-2-HA with GFP-Rab11A (yellow) or TgGRA7 (white) are shown. (C) immunoEM. HeLa cells transfected with ALG-2-HA, infected with parasites for 20 h and fixed and processed for immunoEM with anti-HA (5 nm-gold) and anti-Rab11A (10 nm-gold) antibodies. (a) ALG-HA and Rab11A in the mammalian cell (b) intra-PV structure with Rab11A membranes surrounded by ALG-2-HA (boxed region, with enlargement). (c, d) Additional illustrations of intra-PV Rab11A vesicles surrounded by ALG-2-HA containing membranes. mm, mammalian cell; Go, Golgi; P, parasite; hc, host cell.

### The ESCRT-interacting proteins TgGRA14 and TgGRA64, secreted to the PVM, function synergically in Rab11A vesicle PV internalization

*T. gondii* secretes two dense granule proteins, TgGRA14 and TgGRA64, that physically interact with several ESCRT components, and localize to the PVM and IVN post-secretion (Rivera-Cuevas et al., 2021; Mayoral et al., 2022). We examined the contribution of TgGRA14 and TgGRA64 to the internalization of host Rab vesicles. First, we compared the PV distribution of endogenous CHMP4B for WT parasites and a strain lacking both *gra14* and *gra64*. CHMP4B was detected along the PVM of Δ*gra14*Δ*gra64* parasites similarly to WT (Fig. 9 A), but little to none were observed within the PV, in contrast to WT PV that displayed several CHMP4B puncta on the IVN. This suggests that CHMP4B may associate with the PV independently of TgGRA14 and/or TgGRA64 but that these proteins are most likely involved in CHMP4B-mediated PVM invaginations. Inspection of Δ*gra14*Δ*gra64* ultrastructure did not reveal obvious defects of the PVM or host organelle association (Fig. 9 B, panel a). The PV contains an IVN (Fig. 9 B, panel b; Fig. S4, panel a) but also abnormal fibrous structures appended to IVN tubules (Fig. S4, panels b and c). No organellar abnormalities were observed for Δ*gra14*Δ*gra64* parasites, except for the accumulation of starch-like amylopectin granules that are abundant in *T. gondii* bradyzoites in cysts and barely apparent in fast-replicating tachyzoites unless these are stressed (Coppin et al., 2003). TEM observations of the PV of Δ*gra14* parasites show the same phenotypic features as for Δ*gra14*Δ*gra64*, including the generation of amylopectin stores (Fig. 9 B, panels c and d). Accumulation of amylopectin granules suggest that parasites lacking TgGRA14 and/or TgGRA64 probably undergo remodeling of their metabolic state to cope with stress conditions, such as nutrient limitations.

**Figure 9.**
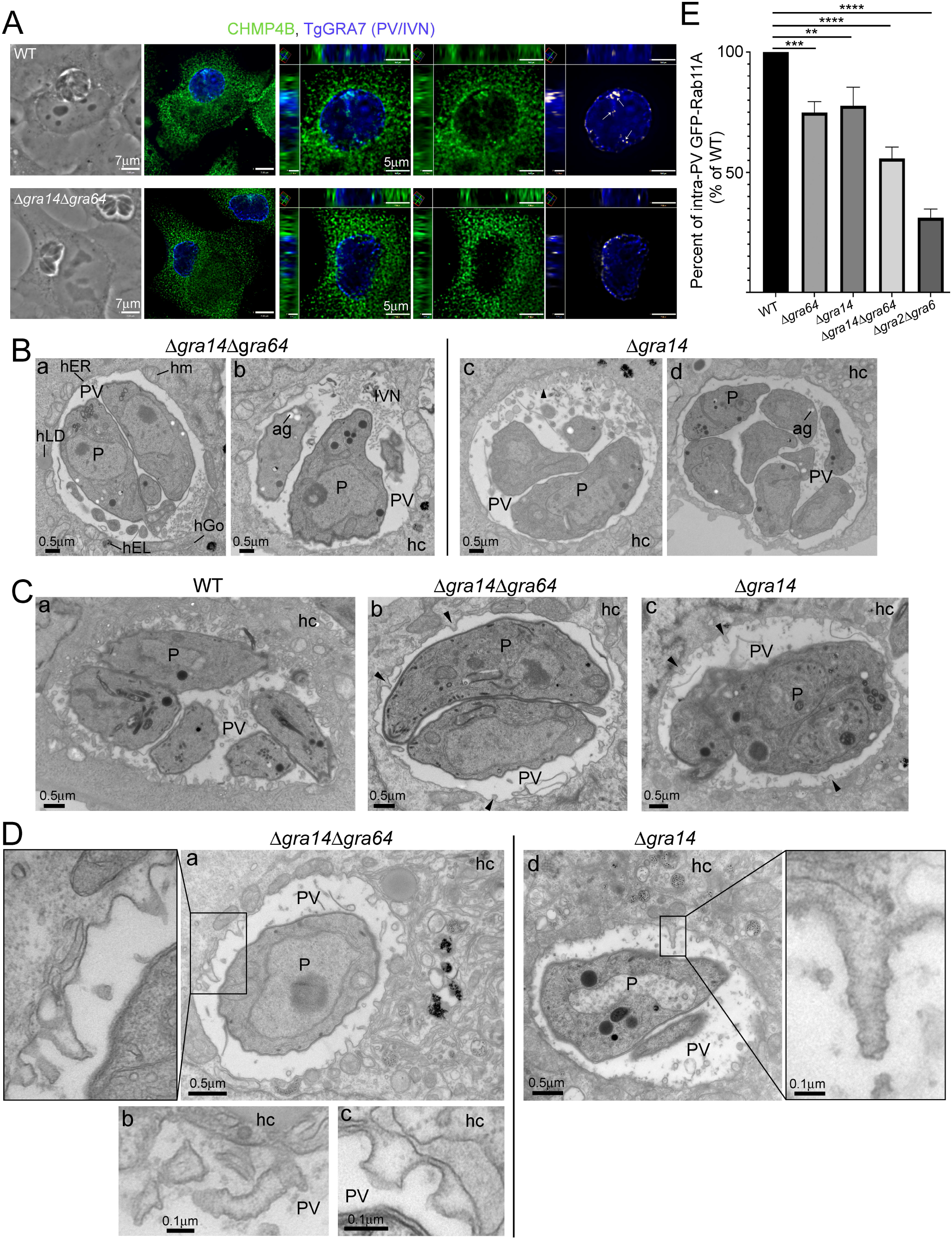
Defects in Δ*gra14*, Δ*gra64* and Δ*gra14*Δ*gra64* parasites in PVM remodeling and Rab11A vesicle internalization. (A) IFA. HeLa cells were infected with WT or Δ*gra14*Δ*gra64* parasites for 20 h, fixed and immunostained for CHMP4B and TgGRA7. Individual z-slices and cropped images of the PV (orthogonal views) are shown plus positive PDM images (white). (B-D) EM. HeLa cells, non-transfected (B) or transfected with CHMP4B-mEm (C, D) were infected with WT, Δ*gra14*Δ*gra64* or Δ*gra14*Δ*gra64* parasites for 20 h and incubated with LDL-gold for ultrastructural analysis. Arrowhead points to unidentified fibrous structure. ag, amylopectin granule; hc, host cell; hEL, host endo-lysosome; hGo, host Golgi; hLD, host lipid droplet; hm, host mitochondrion; P, parasite. (E) VERO cells stably expressing GFP-Rab11A were infected with WT or indicated knockout mutant parasites for 20 h, fixed and immunostained with TgGRA7. The number of intra-PV Rab11A puncta were measured using Volocity. The mean number of intra-PV GFP-Rab11A puncta was calculated for each sample per experiment and converted into percent of WT. Then, the mean ±SD was calculated for the 3 experiments and plotted. Mean values ± SD, *n* = 3 independent experiments, percent of WT is shown. At least 22 PV were measured per sample per experiment. p-values, **=0.0015, ***=0.0006, ****<0.0001

Second, we investigated whether Δ*gra14*Δ*gra64* and Δ*gra14* parasites remodel their PVM to create buds or invaginations in HeLa cells transfected with DN CHMP4B-mEm by EM. While the PVM of WT parasites displayed numerous, long tubular extensions enclosing CHMP4B-mEM filaments on each parasite section viewed (Fig. 9 C, panel a), the PVM of Δ*gra14*Δ*gra64* and Δ*gra14* parasites exhibited sparse buds (Fig. 9 C, panels b-d). In rare sections of either mutant, longer tubular invaginations were observed, containing inner filaments along the tubule membrane and displaying irregular constrictions (Fig. 9 D, panels a-c) or sometimes short ramifications (Fig. S4 B). In 22% of sections examined for Δ*gra14*Δ*gra64* parasites, no PVM budding or bending induced by CHMP4B filaments was apparent, reflecting the rarity of PVM remodeling events in the double mutant (Fig. S4 C).

Lastly, we compared the ability of Δ*gra14*, Δ*gra64*, Δ*gra14*Δ*gra64* and Δ*gra2*Δ*gra6* parasites to scavenge host Rab11A vesicles, relative to WT parasites. VERO expressing GFP-Rab11A were infected with each parasite strain to assess the number, distance from the PV centroid and size of intra-PV GFP-Rab11A puncta. Compared to WT parasites, both Δ*gra14* and Δ*gra64* parasites showed a statistically significant ∼25% decrease in the number of intra-PV host GFP-Rab11A puncta (Fig. 9 E and Fig. S5 A). A more pronounced reduction in puncta number was observed for Δ*gra14*Δ*gra64*, corresponding to a ∼45% decrease compared to WT parasites, though still not as dramatic as for Δ*gra2*Δ*gra6* parasites with a ∼70% decrease. No differences were observed in the distance to the PV centroid nor in the volume of intra-PV GFP-Rab11A puncta between WT and each mutant, except for Δ*gra2*Δ*gra6* (Fig. S5 B and C). Collectively, our data indicate a connection between the poor ability of *Toxoplasma* lacking *gra14* and *gra64* to remodel the PVM and their reduced competency in sequestering intra-PV host vesicles. However, the retention of some activity reveals that other protein/s may be able to recruit host ESCRT at the PVM and function analogously for host vesicle uptake. Our results suggest a synergistic role for TgGRA14 and TgGRA64, which both function to recruit host ESCRT components, in host vesicle uptake. It also emphasizes the foremost contribution of IVN tubules in entrapping host organelles.

## Discussion

For any obligate intracellular pathogen, life within the host intracellular milieu requires major investments to ensure a steady nutrient supply. As such, *Toxoplasma* is proficient in manipulating its host cell to gain access to vital nutrients, either from the cytosol (e.g., molecules, proteins) or from organelles (e.g., lipids). One of the strategies developed by *Toxoplasma* for nutrient import is the formation of inward PVM buds, creating a pathway for entry of host material into the PV. TEM examinations of the PVM illustrate various size and shapes (from vesicular to tubular) of PVM buds. We previously described two types of PVM invaginations that are depicted in our model in Fig. 10 A: one generated by host microtubules recruited at the PV and poking into the PVM (Coppens et al., 2006), and the other mediated by IVN tubules secreted by *Toxoplasma* in the PV and fusing with the PVM (Romano et al., 2017). We also identified smaller PVM buds morphologically distinct from these two tubular PVM invaginations. Host endocytic structures are often observed encroached in these different PVM invaginations before being engaged deeper into the PV lumen. ESCRT proteins remodel membranes by creating indentations and forming a bridge between membranes to bring them close enough for fusion and fission. *Toxoplasma* expresses two proteins (TgGRA14 and TgGRA64) previously shown to interact with ESCRT machinery at the PVM and IVN (Rivera-Cuevas et al., 2021; Mayoral et al., 2022). In this study, we investigated how *Toxoplasma* exploits host ESCRT machinery by focusing on 4 proteins (ESCRT-III component CHMP4B; accessory proteins ALIX and ALG-2; AAA-type ATPase Vps4) for PVM budding and scission in the context of the intra-PV sequestration of host endocytic organelles. On the parasite side, we examined the contribution of the two ESCRT interactors, TgGRA14 and TgGRA64, to facilitate this process. Based on our findings combined with published data, we propose a model in Fig. 10 B summarizing the interplay between these host and parasite proteins for entrapping some host organelles into the PV.

**Figure 10.**
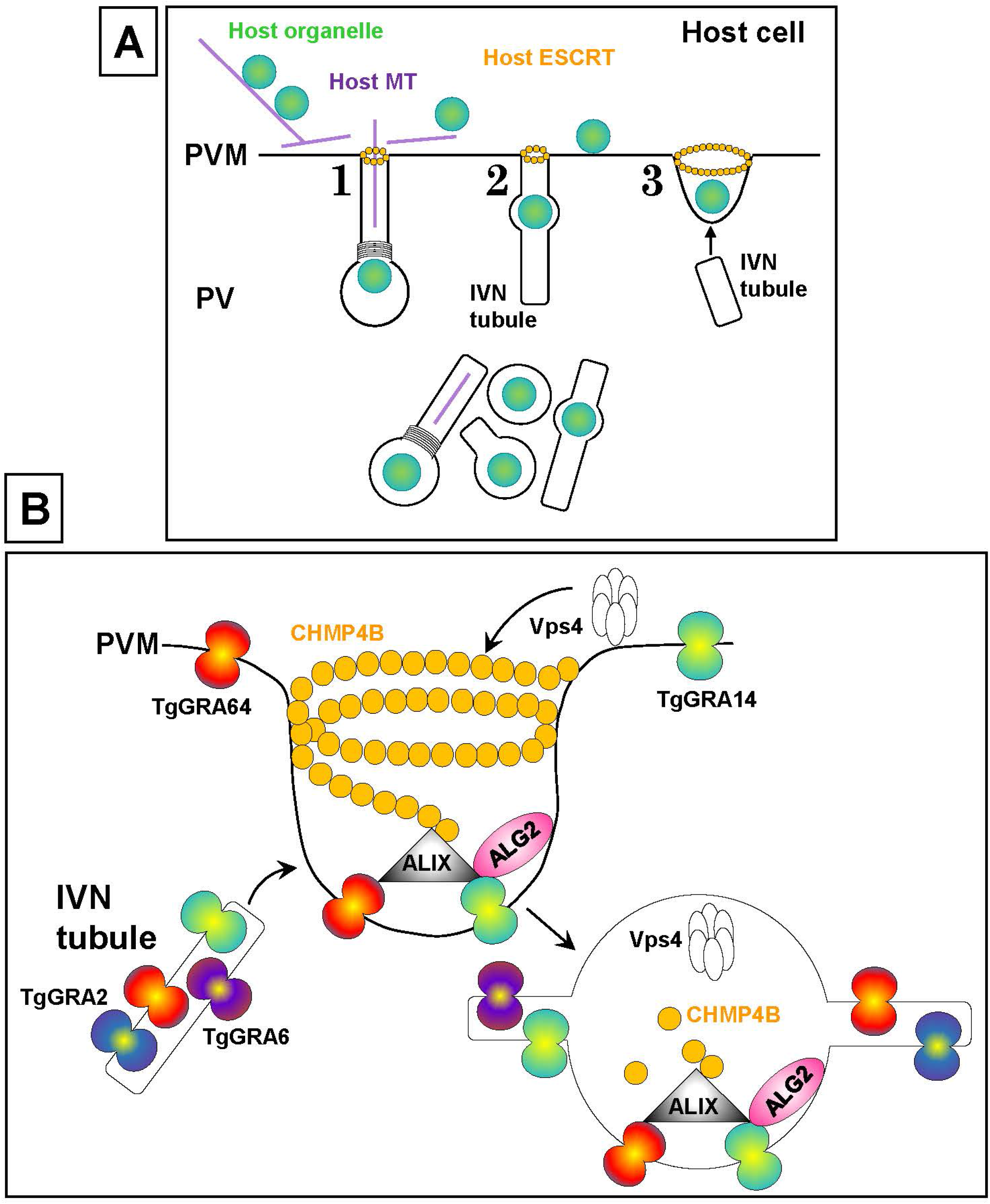
Prospective model of host ESCRT-III components and Vps4 at the PV for internalization of host organelle in synergy with the IVN. (A) Overview of cellular events leading to host organelle trafficking along microtubules and internalization into the PV. Three PVM invaginations are depicted: 1. Host microtubule-based invagination; 2. IVN tubule fused with the PVM; 3. Bud with broad neck. Host ESCRT-III components are recruited at the neck of these invaginations for detachment from the PVM. (B) Molecular players involved host organelle internalization (host organelles not shown for clarity). TgGRA14 and TgGRA64 at the PV bind to ALIX interacting with CHMP4B subunits that cluster, resulting in deep membrane concavities with the walls of the membrane neck close enough for fusion prior to Vps4 scission activity. Concomitant, IVN tubules fuse with CHMP4B-induced tubules, contributing to their elongation and supply in TgGRA14 and TgGRA64. Upon PVM detachment, tubulo-vesicles derived with PVM and IVN containing host organelles and residual ESCRT-III components/Vps4, accumulate inside the PV.

ESCRT proteins govern membrane remodeling processes that require stabilizing membrane curvature, such as the formation of endosomal ILV in MVB. ESCRT machinery is composed of functionally distinct subcomplexes with ESCRT-III and the ATPase Vps4, universally involved in ESCRT-dependent membrane remodeling processes (Vietri et al., 2020; Schöneberg et al., 2017; Pfitzner et al., 2021; Hurley, 2015). Similar to the budding of the limiting membrane of MVB, we propose that ESCRT components recruited to the PV generate PVM invaginations protruding into the PV lumen. By analogy to ILV in MVB, ESCRT-III binds inside the neck of the forming bud to cleave PVM buds away from the host cytosol and PVM-derived vesicles/tubules accumulate in the PV lumen. The formation of ILV derived from the MVB membrane and intra-PV vesicles/tubules from the PVM can then be viewed as topologically comparable. However, aspects of the process would differ especially in terms of ESCRT-III recruitment to the PV that would involve specific parasite proteins. The recruitment of ESCRT-III to distinct membranes is accomplished through compartment-specific targeting factors plus assembly factors. For the formation of ILV in MVB, ESCRT-0, ESCRT-I and ESCRT-II function concurrently to recruit ESCRT-III (Vietri et al., 2020; Schöneberg et al., 2017; Hurley, 2015), while in cytokinetic abscission the centrosomal Cep55 protein recruits TSG101 from ESCRT-I and ALIX to the midbody preceding the recruitment of ESCRT-III (Lee et al., 2008; Elia et al., 2011; Christ et al., 2016). Some viruses have evolved to intercept host ESCRT machinery by secreting virus-specific targeting/assembly factors to facilitate their egress from the host cell. For example, the HIV-1 Gag protein interacts with TSG101 and ALIX, leading to the recruitment of ESCRT-III and Vps4 to the plasma membrane to induce budding of the viruses from the host cell surface (Strack et al., 2003; Pornillos et al., 2003). In the case of *Toxoplasma*, proximity-biotinylation screens reveal the presence of selective host targeting/assembly factors (TSG101, VSP28, VSP37, ALIX, ALG-2) proximally to the PV (Cygan et al., 2021), with ALIX and ALG-2 localized to the PVM/IVN (Cygan et al., 2021; Rivera-Cuevas et al., 2021); A bioinformatics analysis of the *Toxoplasma* genome identified TSG101-and ALIX-targeting motifs in the sequence of TgGRA14 that are exposed to the host cytosol; the Δ*gra14* mutant diminishes PVM recruitment of endogenously expressed GFP-TSG101 but has no effect on ALIX, suggesting the involvement of parasite protein/s other than TgGRA14 for ALIX retention at the PVM (Rivera-Cuevas et al., 2021). Another co-immunoprecipitation and proximity labeling study shows that TgGRA64 interacts with several ESCRT components, including CHMP4B, TSG101, ALG-2, and ALIX (Mayoral et al., 2022). Our immunofluorescence assays show that the Δ*gra14*Δ*gra64* mutant is still able to recruit CHMP4B at the PVM, indicating the expression of other ESCRT recruiter/s that compensate for the loss of *gra14* and *gra64*. This highlights that the attraction of several ESCRT components to the PV to shape (or repair) the vacuolar membrane involves more than 2 parasite proteins and once again demonstrates the versatility of *Toxoplasma* and the redundancy of its protein repertoire to adapt to genetic or environmental changes.

ESCRT-III is a heteropolymer composed of α-helical CHMP proteins, with CHMP4 being the most abundant (Teis et al., 2008). ESCRT-III subunits perform their roles in membrane deformation by changing their conformation and polymerizing into membrane-remodeling filaments. CHMP4A and CHMP4B assemble into filament spirals that attach to membranes, resulting in membrane tubulation. In *Toxoplasma*-infected cells, endogenous CHMP4B is mostly cytosolic in the host cell with partial association with the PVM and IVN, indicating that CHMP4B-containing buds formed at the PVM are subsequently internalized into the PV at sites near the IVN. CHMP4 has multiple domains: an N-terminal membrane-binding “ANCHR” domain, a core domain of four tightly bundled α-helices, a flexible linker, an autoinhibitory α-helix that folds back against the core domain, a MIT-interacting motif MIM that interacts with Vps4, and an α-helix (α6) that interacts with Bro1/ALIX (Tang et al., 2015). The autoinhibitory domain mediates CHMP4 cycling between a closed, inactive conformation diffusely localized to the cytosol and an open, active conformation associated with membranes. The addition of fluorescent tag proteins or epitope tags to either end of CHMP4B, especially coupled with overexpression, leads to DN effects on ESCRT processes, and has been a useful tool to dissect the functions of CHMP4B on the *Toxoplasma* PV. Overexpression in infected cells of CHMP4B fused C-terminally with mEmerald generate impressive tubular invaginations, from the PVM, enclosing CHMP4B spiral filaments. Membrane-associated ESCRT-III usually has a short lifespan (Baumgärtel et al., 2011; Jouvenet et al., 2011) due to the activity of Vps4 in disassembling and recycling ESCRT-III subunits (Babst et al., 1998; Lin et al., 2005). Depletion of Vps4A and B, however, can increase the lifetime of membrane-associated ESCRT-III, leading to the accumulation on endosomal structures and the plasma membrane of ESCRT-III filaments composed of uniform spiral assemblies with a mean outer diameter of 110 nm (Cashikar et al., 2014). Our DN CHMP4B-mEm system mimics a Vps4 depletion condition, with the accumulation of stable PVM invaginations containing very long (up to 7 µm) and narrow (110 nm in diameter) CHMP4B filaments, reflecting a blockage of PVM scission. The stabilization of CHMP4B in its assembled state within PVM invaginations may be due to poor access of Vps4 to the MIM domain of CHMP4B-mEm or insufficient amounts of Vsp4 enzyme to depolymerize all the CHMP4B filaments at the PVM.

The depth of a membrane deformation induced by ESCRT-III filaments is influenced by three factors: filament geometry (depending on the copolymer composition), preferred filament radius (the larger resulting in shorter helices that give rise to shallower conical deformations) and filament stiffness (increasing rigidity results in changes in shape from conical toward tubular) (Harker-Kirschneck et al., 2019). For tubular deformation, the internal filament energy dominates over the elastic energy required to bend the membrane. Observations of multiple and deep tubular invaginations of the PVM upon CHMP4-mEm overexpression suggests the assembly of CHMP4 into very long polymers that form highly rigid filaments with small radii. Another factor that may contribute to long PVM tubule formation could be the density of CHMP4B recruitment by membrane-bound parasite proteins at the PVM that recruit CHMP4B targeting factors (e.g., TgGRA14). For example, TgGRA14 and TgGRA64 may induce PVM invaginations through interactions with ALIX-CHMP4. Indeed, by EM, fewer CHMP4B-mEmerald invaginations are detected in the Δ*gra14* mutant, with even fewer present in Δ*gra14*Δ*gra64*. Uniquely compared to other systems, CHMP4B-associated buds or tubules at the PVM seem to fuse with IVN tubules based on data showing sparse and short PVM invaginations in the Δ*gra2*Δ*gra6* mutant under DN CHMP4B-mEm conditions. Through the fusion of IVN tubules with CHMP4B-associated invaginations, additional membranes are supplied creating deeper invaginations in which host material may remain trapped, with no possible return to the host cell.

Detection of ALIX, ALG-2, CHMP4B and Vps4 on the IVN containing host organelles would entail the participation of these proteins in host organelle internalization into the PV through the initial recruitment of ALIX-ALG-2 to the PVM, the subsequent attraction of CHMP4B to create indentations, and lastly detachment from the PVM by Vps4. Under expression of the DN mutant Vps4A EQ, very large membrane-bound compartments packed with many host endo-lysosomes are observed inside the PV, with a morphology distinct from host endo-lysosomes-containing IVN tubules typically found in PV of WT, indicating that the lack of functional Vps4 interferes with proper internalization. The same phenotype is observed when FLAG-CHMP4B is overexpressed, with these large structures still connected to the PVM, suggesting impaired scission activity. While the Vps4 binding domain is near the C-terminal end of CHMP4B, it remains possible that the acidic FLAG tag interferes with the CHMP4B ANCHR domain that contributes to membrane binding and, in doing so, may impede proper ESCRT-IIIassociation with membranes and thus Vsp4 function in membrane scission.

In summary, this study has highlighted unique features of ESCRT-mediated processes at the *Toxoplasma* PV. First, intravacuolar membranous tubules likely attach to ESCRT-mediated buds to elongate the invaginations at the PVM prior to scission. Second, ESCRT-mediated invaginations can be involved in the internalization of cargos as large as endosomes or Rab vesicles into the PV. Instead, the ESCRT machinery appears to function in a process more akin to phagocytosis for the engulfment of large particles. Lastly, many parasite proteins and perhaps distinct complexes may be involved in interfacing with host ESCRT at the PV. To this point, the Δ*gra14*Δ*gra64* mutant has a greater defect in host Rab11A vesicle internalization and in the formation of CHMP4B-mEmerald induced PVM invaginations than single mutants, suggesting a synergistic role for TgGRA14 and TgGRA64 at the PVM/IVN through interactions with specific ESCRT components.

## Materials and Methods

### Reagents and antibodies

All reagents were obtained from Sigma-Aldrich or Thermo Fisher Scientific, unless otherwise stated. The following primary antibodies were used in this study: rat monoclonal and rabbit polyclonal anti-GRA7 (Coppens et al., 2006). Commercial antibodies include rabbit polyclonal anti-Rab11A (Cell Signaling, Danvers, MA), rabbit monoclonal anti-myc (clone 71D10; Cell Signaling), rabbit polyclonal anti-GFP antibodies (Thermo Fisher Scientific), mouse monoclonal anti-HA.11 (clone 16B12; Biolegend, San Diego, CA), rabbit polyclonal anti-CHMP4B (ProteinTech, Rosemont, IL), and mouse monoclonal anti-FLAG (clone M2; Sigma). Secondary antibodies used were: goat anti-rat, anti-rabbit and anti-mouse conjugated to Alexa Fluor 488, 594, or 350.

### Mammalian cell and parasite culture

Human foreskin fibroblasts (HFF), HeLa and VERO cells were obtained from the American Type Culture Collection (Manassas, VA). VERO stably expressing GFP-Rab11A was described previously (Romano et al., 2017). All cells were grown in alpha-minimum essential medium (Alpha MEM) supplemented with 10% fetal bovine serum (FBS), 2 mM glutamine and penicillin/streptomycin (100 units/ml per 100 µg/ml) and maintained at 37°C in 5% CO_2_. The tachyzoite RH strain (type I lineage) of *Toxoplasma* was used in this study. RH deleted for both *gra2* and *gra6 (Δgra2Δgra6*) was generously provided by M.-F. Cesbron-Delauw (Université Grenoble Alpes, Grenoble, France; (Mercier et al., 2002)). The Δ*gra14*, Δ*gra64* and Δ*gra14*Δ*gra64* strains were described previously. Generation of the RH Δ*gra14* strain was described previously (Rivera-Cuevas et al., 2021). The RH Δ*gra64* strain (made in a Δ*ku80Δhxgprt* background for easier genetic manipulation) was generated identically to the Δ*gra64* strains described in (Mayoral et al., 2022). The Δ*gra14*Δ*gra64* strain was generated from the RH Δ*gra64* strain as follows: first, CRISPR-Cas9 gene editing was used to target the endogenous *GRA14* locus by transfecting 7.5 µg of a vector expressing an HXGPRT resistance cassette and Cas9 with the guide RNA sequence 5’ – CCAGAGACCAAGCGAATAGA – 3’, along with 1.5µg of a repair template generated by PCR (Forward primer: 5’–TCCGAGTTTACACAGGCTGGGCTACCAGAGACCAAGCGAAGAACAGAAACTGATCTCAGA-3’; Reverse primer: 5’-GGTTCCCGATTGGTCTACATTACTGACTTCAACCCTTCTAAAGATCCTCTTCGGAAATAA –3’), designed to insert three copies of the myc epitope-tag at the C-terminus of the gene for the purposes of confirming the absence of endogenous GRA14 protein after gene knockout. Following transfection, parasites were selected for in mycophenolic acid (25 µg/mL) and xanthine (50 µg/mL) for one week. Transfected parasites were subcloned by limiting dilution in 96-well plates. Successful tagging was demonstrated by IFA using anti-myc tag antibody and by Sanger Sequencing the PCR products amplified from genomic DNA, using the following primers to amplify the C-terminus of GRA14: 5’-CGCCGAACCTGAACACAGAT – 3’ and 5’ – ATCACGCGTTCAATCACATC – 3’. After generating a Δ*gra64,GRA14-3xMyc* parasite strain, a CRISPR-Cas9 strategy was again used as described above, transfecting parasites with 7.5 µg of a Cas9 vector containing the guide RNA sequence 5’-GCGCGGGGGGACCGCTCGTC – 3’ (targeting the *GRA14* start codon region) along with 280 pmol unannealed 100bp donor oligos containing homologous arms and multiple stop codons in place of the endogenous GRA14 start codon (5’ – CCACTGCTAAGCACAAAATGCAGGCGATAGCGCGGGGGGTAACTGACTGACTAGCTAACGGGACCGCTCGTCGGGGTGGTCGAGTTGTAGCTGGCTTTTC-3’, and the reverse complement of this sequence). Selection, subcloning, and knockout screening was performed as described above, using the following primers to amplify the 5’UTR and N-terminus region of the GRA14 locus to confirm the insertion of stop codons as designed: 5’-AGAAAGCGTCACTCAGACAT – 3’ and 5’ – CTCCTAGATAGCTTACGGGC – 3’. All parasites were propagated by serial passage in monolayers of HFF (Roos et al., 1994; Khan and Grigg, 2017).

### Plasmids

Plasmids used in this study include mEmerald-Mito-7 (a gift from Dr. Michael Davidson (Addgene plasmid # 54160; http://n2t.net/addgene:54160; RRID:Addgene_54160)), mEmerald-MAP4-C-10 (a gift from Dr. Michael Davidson (Addgene plasmid # 54152; http://n2t.net/addgene:54152; RRID:Addgene_54152)), (Planchon et al., 2011). ALIX-HA and ALG-2-HA (generously provided by Dr. John Boothroyd, Stanford University, Stanford, CA, (Cygan et al., 2021). FLAG-CHMP4B (kindly provided by Dr. Phyllis Hanson, University of Michigan, Ann Arbor, MI), mCherry-N1-VPS4A and mCherry-N1-VPS4A EQ (graciously provided by Dr. Akira Ono, University of Michigan, Ann Arbor, MI (Rivera-Cuevas et al., 2021)) and pCHMP4B-mCherry (generously provided by Jennifer Lippincott-Schwartz, Janelia Research Campus, Ashburn VA (Elia et al., 2011)).

To generate a plasmid with CHMP4B tagged at the 3’end with mEmerald, we amplified CHMP4B from pCHMP4B-mCherry using primers CHMP4B_Cterm forward (5’ ATCCGCTAGCATGTCGGTGTTCGGGAAG 3’) and CHMP4B_C-term reverse (5’ GTGGATCCCCCATGGATCCAGCCCAGTTC 3’) and we amplified mEmerald including vector sequence but omitting the mitochondrial targeting sequence from the mEmerald-Mito-7 plasmid using primers mEmeraldVector forward (5’ TGGATCCATGGGGGATCCACCGGTCGCC 3’) and mEmeraldVector reverse (5’ ACACCGACATGCTAGCGGATCTGACGGTTCAC 3’) and combined the fragments using a HiFi assembly kit (New England Biolabs, Ipswich, MA) to create a plasmid expressing CHMP4B-mEmerald, with a 7aa linker between CHMP4B and mEmerald. To generate a plasmid with CHMP4B tagged at the 5’ end with mEmerald, we amplified CHMP4B from pCHMP4B-mCherry using primers NlinkCHMP4B_fwd (5’ GCTGTACAAGTCCGGACTCAGATCT 3’) and NlinkCHMP4B_rev (5’ CGGTGGATCCTTACATGGATCCAGCCCAG 3’), mEmerald from mEmerald-MAP4-C-10 using primers N_Emerald_fwd (5’ GGTCGCCACCATGGTGAGCAAGGGCGAG 3’) and N_Emerald_rev (5’ TGAGTCCG GACTTGTACAGCTCGTCCATGC 3’), and vector sequence from mEmerald-MAP4-C-10 using mEmMAPVector_fwd (5’ ATCCATGTAAGGATCCACCGGATCTAGATAACTGATCATAATCAGC 3’) and mEmMAPVector_rev (5’ TGCTCACCATGGTGGCGACCGGTAGCGC 3’) and combined the fragments using a HiFi assembly kit to create a plasmid expressing mEmerald-CHMP4B, with a 12aa linker between mEmerald and CHMP4B.

### LDL-gold preparation

Human LDL was isolated from fresh serum by zonal density gradient ultracentrifugation as described (Poumay and Ronveaux-Dupal, 1985). Briefly, the density of human serum was increased to 1.25 g/ml with KBr, centrifuged at 81,900 g x min (Beckman MLA-80 rotor; Beckman Instruments, Inc., Palo Alto, CA) for 24 h at 4°C. The supernatant (lipoproteins) was collected, adjusted to the density of 0.126 g/ml, placed in the bottom of a centrifuge tube, and layered on top with 0.9% NaCl (1:5 vol/vol). After ultracentrifugation at 81,900 g x min for 6 h at 4°C, LDL (middle, yellow layer) was collected, extensively dialyzed against 2 mM sodium borate pH 9, and filtered with a 0.2 µm PVDF syringe filter and stored at 4°C. For the coupling reaction of LDL to gold particles of 15 nm in diameter, a solution of 0.3 mM tetrachloroauric acid was reduced with 1.16mM sodium citrate as described (De Roe et al., 1987) except for the addition of 0.00015% tannic acid for gold stabilization under boiling until the mixture became dark red. For LDL-gold complex preparation, a solution of LDL adjusted at 0.2 mg/ml was added to a solution of gold particles at 57 µg/ml (1:5, vol/vol) for 30 min with gently agitation before addition of 1 ml of 1% BSA in 2 mM sodium borate pH 9. After 5 min, 5 ml of LDL-gold conjugate were layered on 1 ml of a 1.15 M sucrose cushion and sedimented at 97,500 g x min using a Beckman T150 rotor. The solution of LDL-gold conjugates collected from the sucrose solution were dialyzed against PBS before use.

### Transfection and parasite infection

For fluorescence microscopy, HeLa cells were transiently transfected according to the manufacturer’s instructions for JetPrime (Polyplus, Illkirch, France). HeLa cells (50,000) were seeded to coverslips the day before transfection. For the transfection, 0.2 µg DNA in 50 µl JetPrime buffer was mixed with 0.4 µl JetPrime reagent and incubated at room temperature for 10 min prior to addition to cells. VERO cells expressing GFP-Rab11A (40,000) were seeded to coverslips the day before transfection. For the transfection, 0.8 µg of DNA in 50 µl JetOptimus buffer (Polyplus) was mixed with 0.75 µl JetOptimus reagent and incubated at room temperature for 10 min prior to addition to the cells. For EM, HeLa cells were grown in 6-well plates to 60% confluency and transiently transfected with plasmid DNA (1 µg per well) mixed with JetOptimus reagent (1 µl per well) and incubated at room temperature for 10 min, according to manufacturer’s instructions (Polyplus) before adding to cells. All transfected cells were incubated at 37°C with 5% CO_2_ for 4 h, washed with PBS and allowed to recover for 1 h at 37°C with 5% CO_2_ prior to infection with parasites. For infection, freshly egressed parasites were added to the cells for 1 h, washed with PBS to remove extracellular parasites and incubated for times indicated in Alpha MEM. For some samples, LDL complexed with gold was added in Alpha MEM medium with 10% (vol/vol) of delipidated FBS (Cocalico Biologicals, Stevens, PA) and the cells were incubated at 37°C in 5% CO_2_ for 20 h. Infected cells were fixed for either fluorescence microscopy or EM.

### Cell Labeling

For immunolabeling, cells were fixed in PBS with 4% formaldehyde (Polysciences, Inc.) and 0.02% glutaraldehyde for 15 min. Cells were permeabilized with 0.3% Triton X-100 in PBS for 5 min and washed twice with PBS before blocking, except for cells immunolabeled with anti-CHMP4B antibody; for these cells, 0.1% saponin was added to the blocking and antibody buffers and treatment with Triton X-100 was omitted. Cells were incubated in blocking buffer (3% BSA, Fraction V; Thermo Fisher Scientific) in PBS for 1 h followed by incubation in primary antibodies diluted in blocking buffer (1:800 for anti-GRA7; 1:1000 for anti-HA; 1:1500 for anti-FLAG; 1:100 for anti-Rab11A; 1:500 for anti-CHMP4B) for 1 h to overnight. Cells were washed three times with PBS for 5 min each and then incubated in secondary antibody diluted in blocking buffer for 45 min to 1 h followed by three washes with PBS for 5 min each. In some cases, cells were incubated in a 1:1000 dilution of 1 mg/ml DAPI (Roche Diagnostics) in PBS for 5 min followed by three washes with PBS. Coverslips were rinsed with water and mounted on slides with ProLong Diamond mounting solution.

### Fluorescence microscopy

Fixed samples were viewed with a Zeiss AxioImager M2 fluorescence microscope equipped with an oil-immersion Zeiss plan Apo 100x/NA1.4 objective and a Hamamatsu ORCA-R2 camera. Optical z-sections with 0.2 µm spacing were acquired using Volocity software (Quorum Technologies, Puslinch, ON, Canada).

### Image analysis

Images were deconvolved with an iterative restoration algorithm using calculated point spread functions and a confidence limit of 100% and iteration limit of 30-35 using Volocity software. Images were cropped and adjusted for brightness and contrast using Volocity software. Figures were compiled in Adobe Illustrator.

To characterize the intra-PV GFP-Rab11A foci, we acquired optical z-sections of infected cells with PV containing 8 parasites, to normalize for PV size. After deconvolution, images were analyzed with a measurement protocol generated in Volocity (Quorum Technologies) according to published protocols with some modifications (Romano et al., 2017, 2021). The measurement protocol measured objects in 3D reconstructed volumes of optical z-slices. The PV was identified by the fluorescence intensity of TgGRA7 signal; thresholds were set manually by using the outer TgGRA7 PVM staining as the boundary of the PV and the minimum object size was set to 20 µm^3^. To close any holes in the identified structure, the close function was used with 20 iterations. GFP-Rab11A puncta were identified by fluorescence intensity (automatic thresholds with an offset threshold ranging from 110 – 170%, minimum object size of 0.4 µm^3^), noise was removed using a medium filter, and objects were separated using the “separate touching objects” tool with an object size guide of 0.1 µm^3^. Lastly, the compartmentalize function was used to identify GFP-Rab11A objects inside the PV. The number and volume of GFP-Rab11A puncta inside the PV were calculated plus the distance of the intra-PV GFP-Rab11A puncta to the PV centroid.

Mander’s correlation coefficients (MCC) were calculated using Volocity software (Quorum Technologies) along with a product of the difference of the means image (PDM), which is generated by identifying voxels where each signal is above its mean value after thresholding for background. Thresholds were set by measuring, using an ROI, the fluorescence intensity of each channel in a region of the nucleus without staining for either signal (background). The positive PDM shows voxels where both fluorescence signals are above the mean and show positive correlation.

### Transmission Electron Microscopy (TEM)

For thin-section TEM, cells were fixed in 2.5% glutaraldehyde (Electron Microscopy Sciences; EMS, Hatfield, PA) in 0.1 M sodium cacodylate buffer (pH 7.4) for 1 h at room temperature, and processed as described (Fölsch et al., 2001).

For immunogold staining, cells were fixed in 4% paraformaldehyde (Electron Microscopy Sciences) in 0.25 M HEPES (pH7.4) for 1 h at room temperature, and then in 8% paraformaldehyde in the same buffer overnight at 4oC. They were infiltrated, frozen and sectioned as described (Romano et al., 2017). The sections were immunolabeled with antibodies against GFP at 1:10, HA at 1:50 or Rab11A at 1:10 diluted in PBS/1% fish skin gelatin, and then with secondary IgG antibodies coupled to 10 nm protein A-gold particles before examination with a Philips CM120 EM or a Hitachi 7600 EM under 80 kV, equipped with a dual AMT CCD camera system. The AMT v6.1 software was used for quantitative measurement of the diameter of PVM invaginations with CHMP4B-mEm (from 62 invaginations from 26 different PV sections) and of intra-PV host organelle size and distance performed on 19 to 23 representative PV sections.

### Statistical analysis

Numerical data are presented as means ± SD or in dot plots with means indicated (GraphPad Prism, San Diego, CA). To compare samples, we used a one-way ANOVA with a Tukey’s multiple comparisons test or unpaired two-tailed t-tests using GraphPad Prism.

## Supplemental material

Fig. S1 shows additional examples of CHMP4B tagged N-terminally or C-terminally with mEmerald associating with the PV plus progressive association of CHMP4B-mEm with the PVM. Fig. S2 shows microscopy images and quantification of PV seemingly devoid of parasites in cells expressing CHMP4B-mEm. Fig. S3 shows additional examples of FLAG-CHMP4B associating with the PV and overlapping with intra-PV Rab11A puncta. Fig. S4 details the ultrastructure of Δ*gra14*Δ*gra64* parasites. Fig. S5 the number, distance from the PV centroid and volume of intra-PV Rab11A puncta for parasite mutants lacking *gra14* and/or *gra64*.

## Acknowledgements

We thank the members of the Coppens’ laboratory for helpful discussion during the course of this work and especially Karen Ehrenman for her help in cloning the mEmerald-CHMP4B and CHMP4B-mEmerald plasmids. We are grateful to Phyllis Hanson for helpful advice and discussions related to this work. We also thank the generous providers of plasmids and parasite strains used in this study. We thank the excellent technical staff from Electron Microscopy Core facilities at Johns Hopkins University School of Medicine (M. Delannoy, B. Smith) and Yale University School of Medicine (K. Zichichi). This study was supported by a grant from the NIH R01 AI060767 to I.C and NIH RO1 AI134753 to LMW.

The authors declare no competing financial interests.

## Author Contributions

Conceptualization: JDR, IC, JM, LMW, VBC

Data curation: JDR, IC

Formal analysis: JDR, IC

Funding acquisition: IC

Investigation: IC, JDR

Methodology: JDR, IC, JM, RBG

Resources: JDR, IC, JM, RBG, YR

Validation: JDR, IC

Visualization: JDR, IC

Writing – original draft: JDR, IC

Writing – review & editing: JDR, IC, JM, RBG, YR, LMW, VBC

## Main abbreviations

ALG-2: apoptosis linked gene 2
ALIX: ALG-2-interacting protein X
CHMP: charged multivesicular body protein
CHMP4B-mEm: CHMP4B tagged with C-terminal mEmerald
DN: dominant negative
ESCRT: endosomal sorting complex required for transport
IFA: immunofluorescence assay
IVN: intravacuolar network
mEm-CHMP4B: CHMP4B tagged with N-terminal mEmerald
p.i.: post-infection
PV: parasitophorous vacuole
PVM: PV membrane
TgGRA: *Toxoplasma* dense granule protein
TSG101: tumor susceptibility gene 101 protein
WT: wild-type.

## Supporting information captions

**Figure S1.**
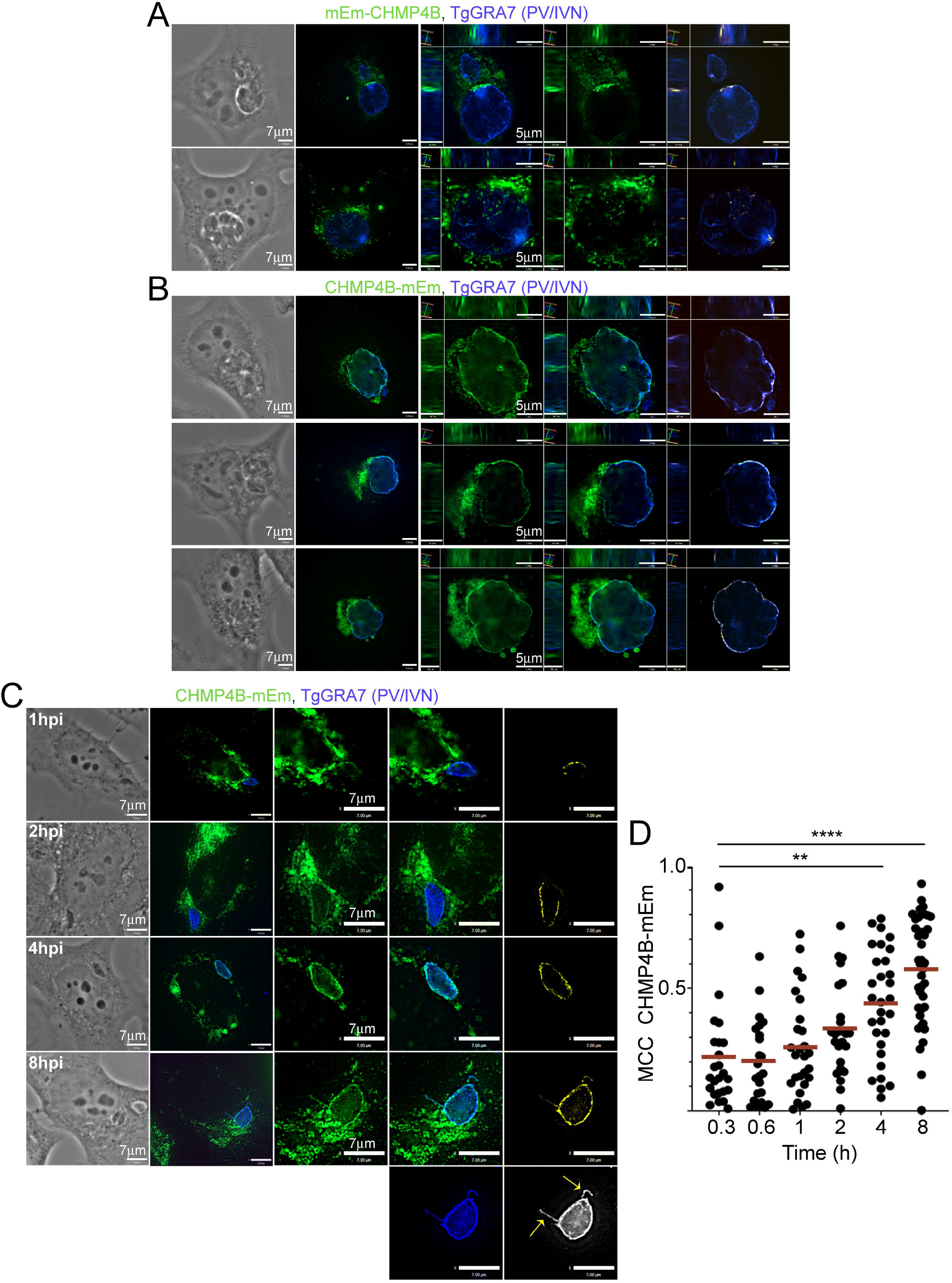
Differential PV localization of mEmerald-CHMP4B and CHMP4B-mEmerald. (A-B) IFA. HeLa cells were transiently transfected with either mEm-CHMP4B (A) or CHMP4B-mEm (B), infected with WT parasites for 20 h, fixed and immunostained for TgGRA7. Individual z-slices and cropped images of the PV (orthogonal views) are shown. A positive PDM image is shown, highlighting in yellow of voxels where both CHMP4B and TgGRA7 intensity values were above their respective means. (C) IFA. HeLa cells transiently transfected with CHMP4B-mEm were infected with WT parasites for the times shown, fixed and immunostained for TgGRA7. A positive PDM image is shown (yellow). Arrows show PVMP containing CHMP4B. (D) HeLa cells transiently transfected with CHMP4B-mEm were infected with WT parasites for the times shown, fixed and immunostained for TgGRA7. Mander’s correlation coefficients (MCC, CHMP4B-mEm) were calculated with Volocity. Dot plots are shown with red bars indicating the mean. At least 20 PV were measured per time point. p-values, ****<0.0001, **=0.0019.

**Figure S2.**
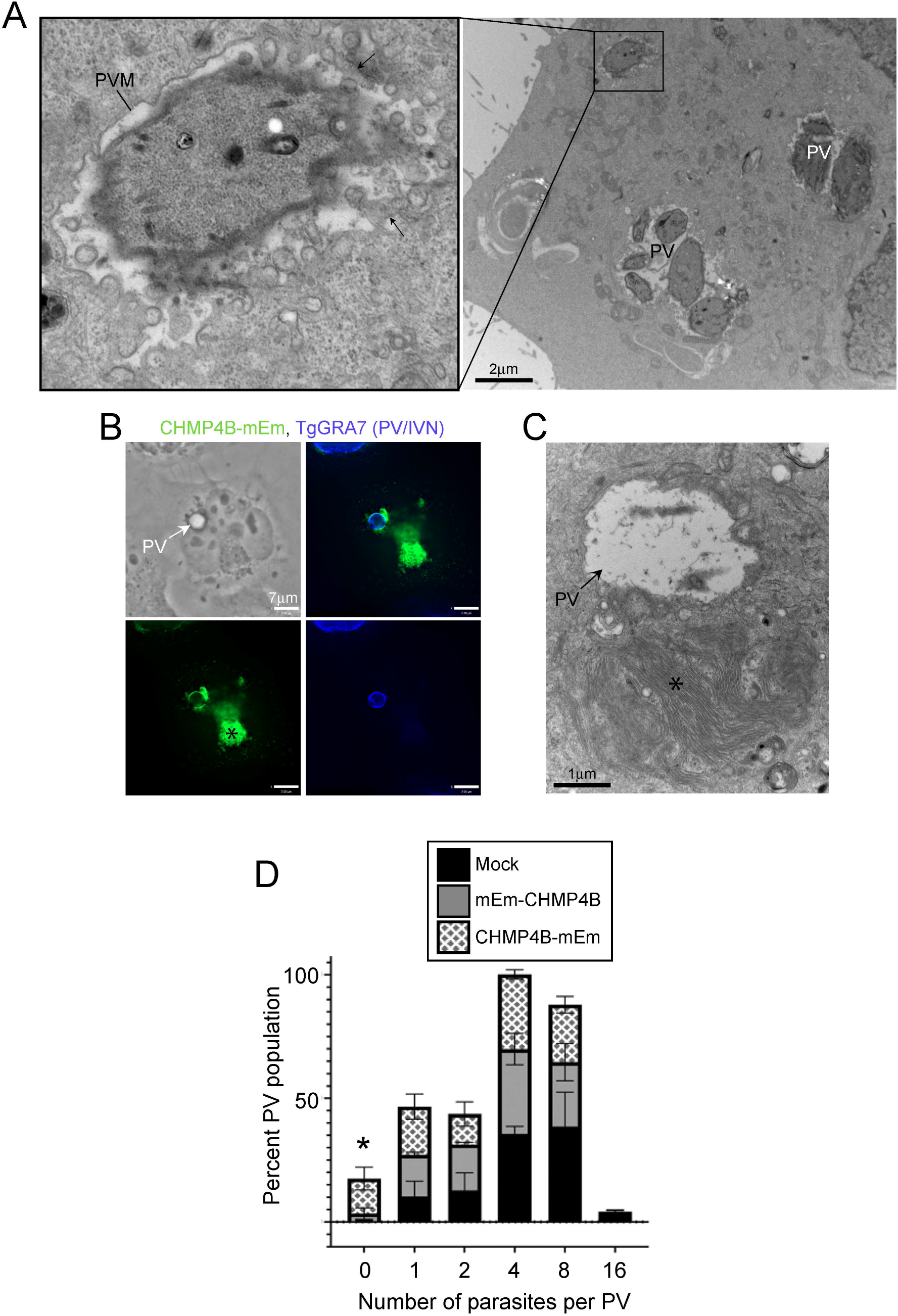
Defective PV in cells expressing CHMP4B-mEmerald. (A) EM. HeLa cells transfected with CHMP4B-mEm and infected with WT parasites for 20 h, showing PV with disorganized parasites in PV, almost no host ER and mitochondria associated with the PVM and many small PV with numerous PVM indentations. (B) IFA. HeLa cells transfected with CHMP4B-mEm were infected with WT parasites for 20 h, fixed and immunostained with anti-GFP and TgGRA7 antibodies. Asterisk shows ‘class E-like’ compartments and arrow a PV devoid of parasite. (C) EM. HeLa cells transfected with CHMP4B-mEm and infected with WT parasites for 20 h, showing an “empty” PV reminiscent of the IFA image in B. (D) Enumeration of parasites within each PV in HeLa cells either mock-transfected or transfected with either mEm-CHMP4B or CHMP4B-mEm and infected with WT parasites for 20 h. Number of parasites equaling zero corresponds to empty PV as in (B). Means ± SD, *n*=3 independent experiments. At least 150 PV (from transfected cells) were counted per sample and PV were defined as positive for TgGRA7 staining. p-values, p=0.0030 (Mock vs. C-tag, 0/PV); p=0.0095 (N-tag vs. C-tag, 0/PV)

**Figure S3.**
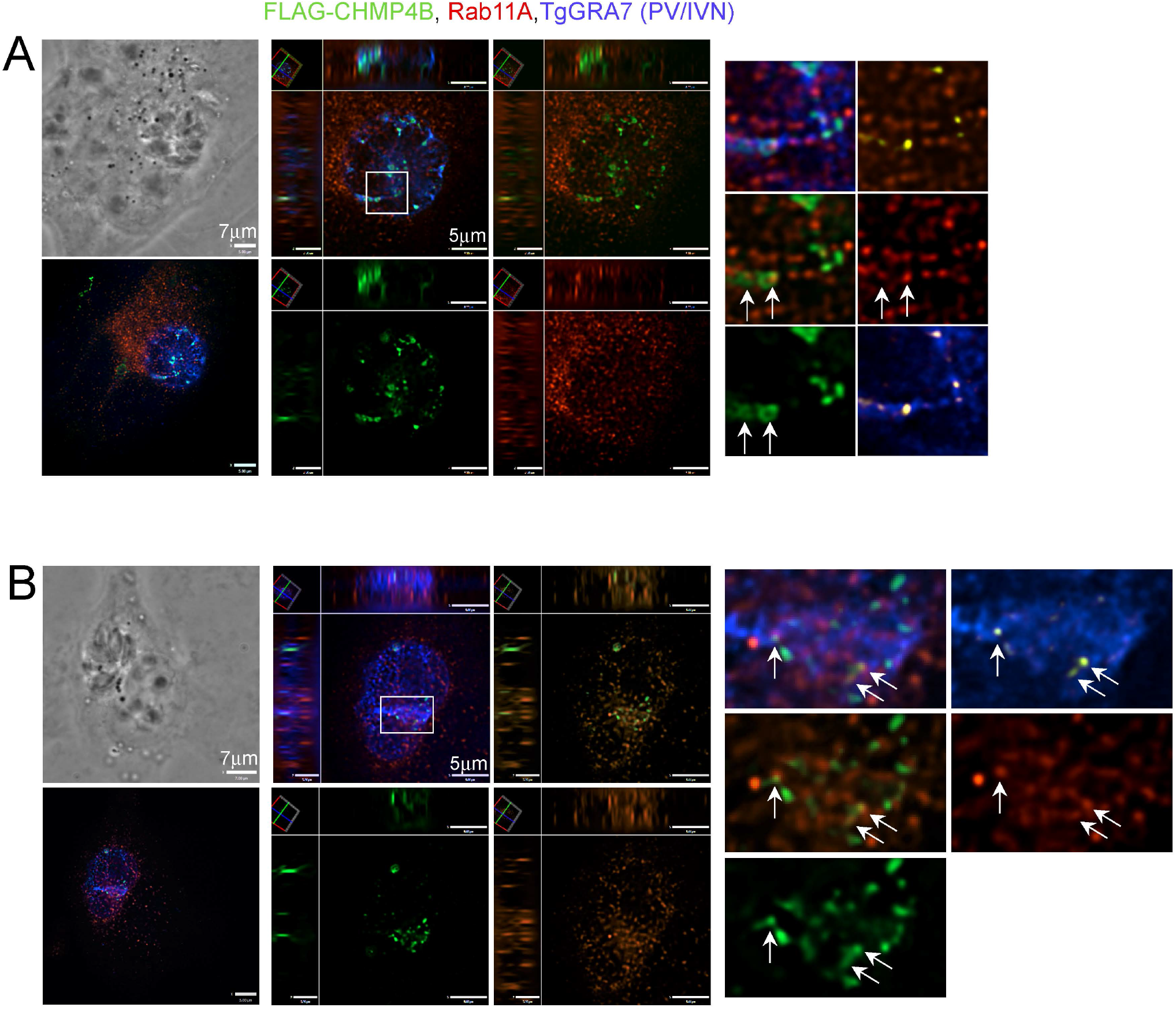
Distribution of intra-PV FLAG-CHMP4B and Rab11A puncta. (A-B) HeLa cells transiently transfected with FLAG-CHMP4B and infected with WT parasites for 20 h were fixed and immunostained for FLAG, Rab11A and TgGRA7. Individual z-slices and cropped images of the PV (orthogonal views) are shown. Many FLAG-CHMP4B puncta associated with the PV (TgGRA7) and some puncta overlap with Rab11A (enlargements, arrows).

**Figure S4.**
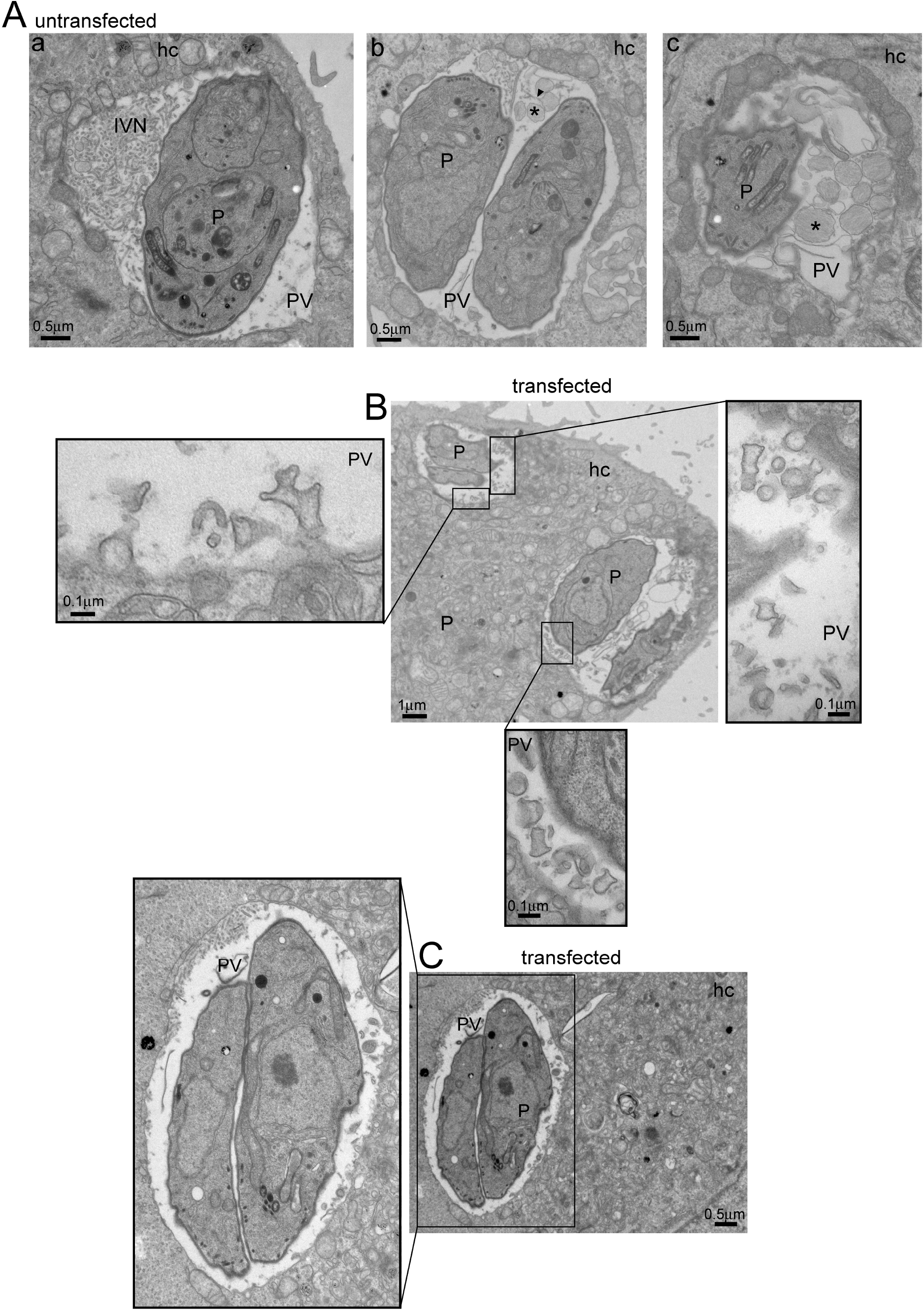
Ultrastructure of Δ*gra14*Δ*gra64* parasites. (A-C) EM. HeLa cells, non-transfected (A) or transfected with CHMP4B-mEmrald (B, C) were infected with Δ*gra14*Δ*gra64* parasites for 20 h and incubated with LDL-gold for ultrastructural analysis. Asterisks point to unidentified fibrous structure. hc, host cell.

**Figure S5.**
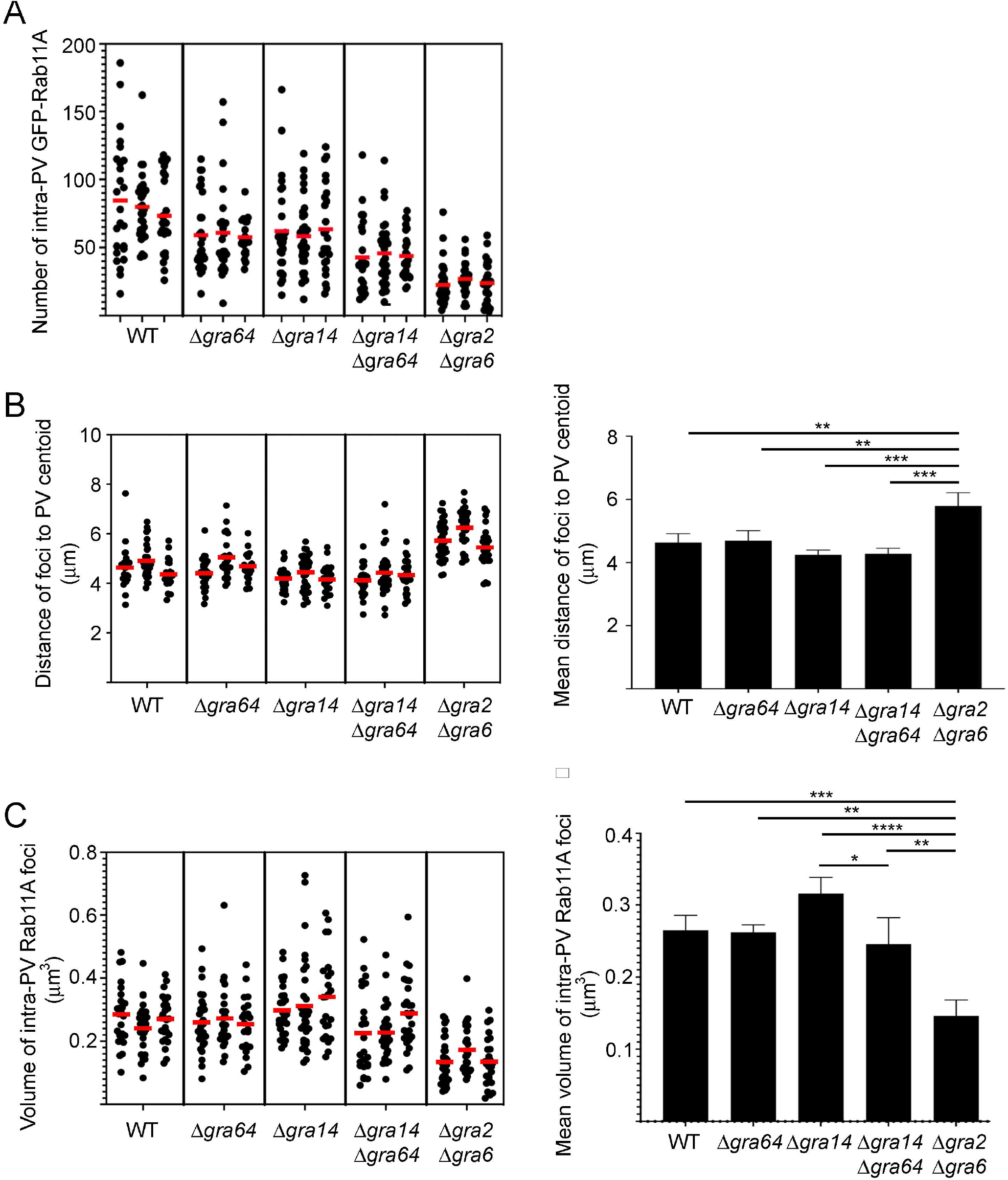
Characteristics of intra-PV GFP-Rab11A puncta for WT, Δ*gra64*, Δ*gra14*, Δ*gra14*Δ*gra64* or Δ*gra2*Δ*gra6* parasites. VERO cells stably expressing GFP-Rab11A were infected with WT, Δ*gra64*, Δ*gra14*, Δ*gra14*Δ*gra64* or Δ*gra2*Δ*gra6* parasites for 20 h, fixed and immunostained with TgGRA7. The number (A), distance from the PV centroid (B) and volume (C) of intra-PV Rab11A puncta were measured using Volocity. At least 22 PV were measured per sample per experiment. (A) Dot plot showing the number of intra-PV GFP-Rab11A puncta for each PV measured for three independent experiments. Red bar, mean. This is the raw data of the mean numbers shown in Fig. 9 E. (B) Dot plot showing the mean distance of intra-PV GFP-Rab11A puncta from the PV centroid for each PV measured for three independent experiments. Red bar, mean. Graph of the mean distance ± SD, *n* = 3 independent experiments. p-values, *** <0.0005, ** = 0.0066. (C) Dot plot showing the mean volume of intra-PV GFP-Rab11A puncta for each PV measured for three independent experiments. Red bar, mean. Graph of mean volume ± SD. *n* = 3 independent experiments. p-values, ****<0.0001, ***=0.0009, **=0.0035, *=0.033.

